# Sequence of the SARS-CoV-2 spike transmembrane domain makes it inherently dynamic

**DOI:** 10.1101/2021.06.07.447334

**Authors:** Sahil Lall, Padmanabhan Balaram, M.K. Mathew, Shachi Gosavi

**Affiliations:** National Centre for Biological Sciences, Tata Institute of Fundamental Research, Bangalore-560065, India; Simons Centre for the Study of Living Machines, National Centre for Biological Sciences, Tata Institute of Fundamental Research, Bangalore-560065, India

**Keywords:** Viral fusion, SARS-CoV-2 spike protein, multiscale molecular dynamics simulations, transmembrane helix self-assembly, single-pass transmembrane helix

## Abstract

The homotrimeric SARS-CoV-2 spike protein enables viral infection by mediating the fusion of the viral envelope with the host membrane. The spike protein is anchored to the SARS-CoV-2 envelope by its transmembrane domain (TMD), which is composed of three TM helices, each contributed by one of the protomers of the homotrimeric spike. Although the TMD is important for SARS-CoV-2 viral fusion and is well-conserved across the Coronaviridae family, it is unclear whether it is a passive anchor of the spike or actively promotes viral fusion. Specifically, the nature of the TMD dynamics and how these dynamics couple to the large pre- to post-fusion conformational transition of the spike ectomembrane domains remains unknown. Here, we computationally study the SARS-CoV-2 spike TMD in both homogenous POPC and cholesterol containing membranes to characterize its structure, dynamics, and self-assembly. Different tools identify distinct segments of the spike sequence as its TM helix. Atomistic simulations of a spike protomer segment that includes the superset of the TM helix predictions show that the membrane-embedded TM sequence bobs, tilts and gains and loses helicity at the membrane edges. Coarse-grained multimerization simulations using representative TM helix structures from the atomistic simulations exhibit diverse trimer populations whose architecture depends on the structure of the TM helix protomer. Multiple overlapping and conflicting dimerization interfaces stabilized these trimeric populations. An asymmetric conformation is populated in addition to a symmetric conformation and several in-between trimeric conformations. While the symmetric conformation reflects the symmetry of the resting spike, the asymmetric TMD conformation could promote viral membrane fusion through the stabilization of a fusion intermediate. Together, our simulations demonstrate that the SARS-CoV-2 spike TM anchor sequence is inherently dynamic, trimerization does not abrogate these dynamics and the various observed TMD conformations may enable viral fusion.

## INTRODUCTION

The cellular infection of the COVID-19 causing SARS-CoV-2 (severe acute respiratory syndrome coronavirus 2) begins by the fusion of the viral envelope with the host cell membrane (*1, 2*) (Fig. 1A). All coronaviruses, including SARS-CoV-2, SARS-CoV and MERS-CoV (Middle East respiratory syndrome coronavirus), rely on envelope-anchored homotrimeric spike proteins to fuse with host cells (*1–3*). The spike protein or simply, spike, is a class I fusion protein which serves both to bind cognate host receptors and to fuse with the host cells (*3, 4*). Each protomer of the SARS-CoV-2 spike homotrimer can be divided into two subunits: the N-terminal S1 that dissociates once the spike has recognized and bound to its host receptor, and the C-terminal S2, which comprises the fusion machinery (*5, 6*) (Fig. 1). Comparison of the pre-fusion (*5, 7–10*) and post-fusion (*6, 11–13*) structures of the SARS-CoV-2 spike protein imply large dynamics of the S2 domain of the spike (Fig. 1A). These S2 dynamics should be conserved across the studied coronaviruses, since the spike structures from different CoVs are highly similar (*5, 9, 10, 14–16*). Furthermore, these dynamics are important because blocking the structural reorganization of the SARS-CoV-2 spike S2 domain can inhibit viral fusion (*17–19*).

**Figure 1:**
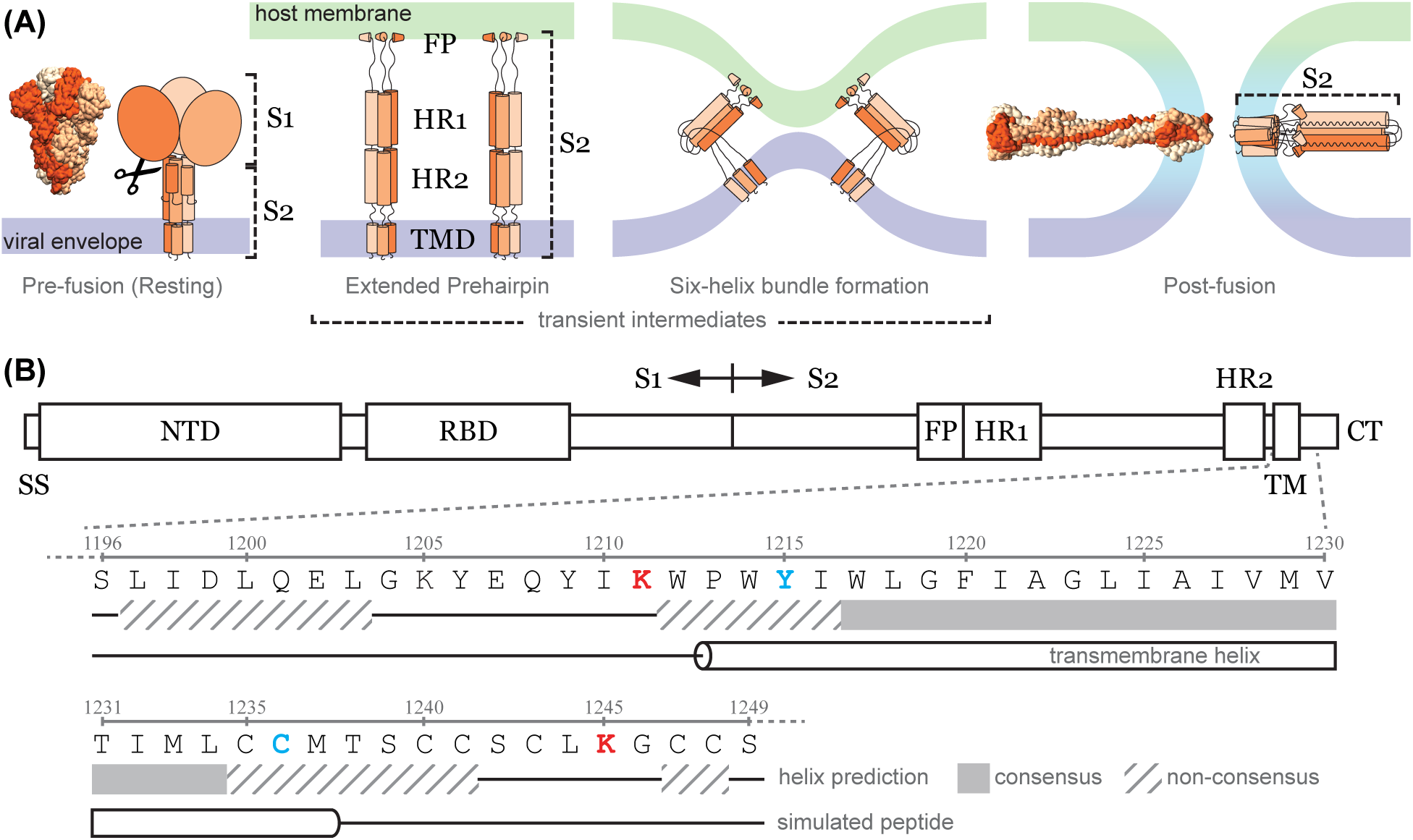
A schematic representation of viral fusion mediated by the Coronavirus spike protein. **(A)** Three transmembrane (TM) helices (TM domain), each contributed by one of the three protomers (shades of orange) of the trimeric spike protein, anchor it to the viral envelope (purple) in the pre-fusion conformation. A protease (scissors) cleavage after the S1 domain frees the S2 to undergo a conformational change that directs the fusion peptide (FP) towards the host membrane (mint) while the TM helices remain anchored to the viral envelope. The two heptad repeat coiled-coil domains (HR1 and HR2) are shown. This intermediate folds upon itself leading to the formation of a six-helix bundle, which brings close the two distal membranes. The endpoint of this is the merger of the two membranes (cyan) and the transition of the spike protein into the post-fusion conformation putting the FP and TM helices in proximity. The available pre-fusion structures of the spike protein, represented here by PDB: 6VXX in the conformation labelled Pre-fusion (Resting) end before the HR2 domain. The complete post-fusion structure (PDB: 8FDW) is shown here in the conformation labelled Post-fusion. **(B)** Simplified domain organization of the spike protein showing the major domains of the spike protein. SS: signal sequence, NTD: N-terminal domain, RBD: receptor binding domain, FP: fusion peptide, HR: heptad-repeat domain, TM: transmembrane helix, CT: C-terminal tail. A section around the TM was used in the atomistic simulations. The sequence of this 54 residue peptide is expanded from the domain organization with two Lys residues (red) marking the ends of the 33-residue long potential TM helix. The results from the various secondary structure and transmembrane domain prediction algorithms (see Fig. S1A) and the peptide segment built as the helix in the atomistic simulations are shown below the sequence. An expanded stretch which includes two additional residues on either side of the consensus peptide is used for analysis (Y1215 to C1236, in cyan).

Several mutational studies of the CoV spike transmembrane (TM) region have established the importance of the correct sequences for optimal infection of host cells (*20–24*). The TM helix is an integral part of the S2 domain and it anchors the spike fusion machinery to the viral envelope (Fig. 1A). Three TM helices, one contributed by each protomeric chain of the homotrimeric spike, together form the TM domain (TMD) of the spike. It is known that the trimeric TMD is required for maintaining the structural integrity of the trimeric pre-fusion state of the coronavirus spike protein (*25*), including that of the SARS-CoV-2 spike (*26*). Regardless, structural information of the spike pre-fusion state comes from water-solubilized spike protein constructs engineered to stay trimeric without the TMD (*5, 7, 9, 27*) (Fig. 1A).

A 20-residue sequence assumed to be the SARS-CoV-2 TM helix forms a stable trimer in solution (*28*), likely reflecting the symmetry of the pre-fusion state, however, several mutations were needed to enable its experimental characterization (*28*). In contrast, the structure of the spike S2 in the post-fusion state (Fig. 1A) reveals no direct interaction between the TM helices (*29*). Together, these observations indicate that there exist assembly and disassembly conformational dynamics between the TM helices in the spike protein. However, without experimentally-derived complete pre-fusion structures of the SARS-CoV-2 spike protein anchored to its viral membrane by the TMD there is a gap in our understanding of the conformational transitions of the TMD and the crosstalk between these dynamics and those of the spike ectomembrane domains (Fig. 1A). Specifically, it is unclear if the TMD is a passive anchor as is implicitly assumed in the design of many experiments (*30, 31*) and simulations (*32, 33*), or if the TMD dynamics facilitate the pre-fusion to post-fusion structural transition of the spike S2 and viral fusion.

Experimental structural information of the TMD of the well-studied human immunodeficiency virus (HIV) fusion protein, gp41 (*34, 35*), and the influenza fusion protein, hemagglutinin (HA) (*36*), were long preceded by molecular dynamics (MD) simulations which indicated their TMD conformations and dynamics (*37–39*). However, most MD simulations of the pre-fusion SARS-CoV-2 spike assume a preliminary trimeric model of TM helices designed to anchor the spike to the envelope (*40, 41*) and only focus on the dynamics of the soluble receptor binding domain (RBD; part of S1) with its ‘up’ and ‘down’ conformations (*42*). Even simulations that focused on conformations of the S2 domain during viral fusion (*33*) restrained the TMD motion during the pre- to post-fusion structural transition. Overall, a lack of knowledge about the conformations and dynamics of the TMD causes it to be schematically represented in models of coronavirus fusion (*1, 2, 43*).

Here, in order to elucidate the structure, organization and self-assembly of the SARS-CoV-2 spike TMD, we first set out to identify the membrane-embedded part of the S2 domain that forms the TM helix. Even that was difficult to identify, because various TM helix prediction algorithms came up with widely varying boundaries for the membrane spanning helix within the S2. So, we performed atomistic MD simulations of an extended segment of the putative TM region from a single spike protomer in a model bilayer. We found that the membrane boundaries of the TM helix in the simulated segment are not constant, and concomitantly the TM helix bobs and tilts in the membrane. We extracted different conformations of the TM helix from these atomistic simulations in order to understand its self-assembly in coarse-grained (Martini) trimerization simulations. We found that different TM helix conformations had different overlapping and at-times conflicting homodimerization interfaces based on whether a series of N-terminal aromatic residues were present (membrane embedded) in the particular TM helix. Combinations of dimeric interface interactions gave rise to two main kinds of trimers in our Martini simulations: symmetric trimers that reflect the symmetry of the pre-fusion state of the spike and asymmetric trimers that may enable the pre- to post-fusion conformational transition of the spike. Our multiscale simulations indicate that both the spike protomer TM helix and its assembly into the spike TMD are inherently dynamic. We discuss these dynamics in the context of viral fusion and the design of molecules that may inhibit fusion by modulating these dynamics.

## RESULTS

### Determination of the membrane-embedded helical fragment of the spike protein

Little structural information exists for either the transmembrane (TM) or the juxtamembrane region of the pre-fusion state of the SARS-CoV-2 spike. This required us to first determine the membrane-embedded residues of a spike protomer. Membrane-spanning helices are usually composed of 21-23 predominantly hydrophobic residues flanked by charged residues (*44, 45*). However, the putative TM region at the C-terminus of the spike protomer has 33 residues flanked by two Lys residues, Lys1211 and Lys1245 that could potentially form the spike TM helix (Fig. 1B). Not just the length, but even the sequence of this 33 residue putative TM region is unusual, with four of the 33 residues being Trp and Tyr (Fig. 1B). The three Trp residues (Trp1212, Trp1214 and Trp1217) along with the Tyr1215 have remained conserved across the relatively benign CoVs discovered in 1960s i.e., the 229E and OC43, as well as the causative agent of the deadly COVID-19: SARS-CoV-2 and its evolved variants of concern. In fact, the entire N-terminal aromatic stretch comprising the three Trp residues, and the Tyr1215 is extremely well conserved across the Coronaviridae family (*20–22, 24*). Here, we term this six residue (Trp1212 to Trp1217) N-terminal fragment of the putative TM helix of the spike protomer the tryptophan-rich aromatic conserved stretch or TRACS.

Furthermore, two Ser, two Thr, three Met and five Cys residues are scattered at the C-terminus of the putative TM fragment (Fig. 1B). Because of this unusual sequence composition, several algorithms that predict TM helices do not converge on consensus start and end residues for the TM helix in the 33-residue long putative TM segment (Figs. 1B and S1A). In order to ascertain the membrane-embedded region of the SARS-CoV-2 spike protomer, we performed atomistic molecular dynamics (MD) simulations using the CHARMM36 force field (Table S1, Methods for details). One copy of a 54 amino acid long polypeptide (Ser1196-Ser1249) was simulated in a homogenous POPC lipid bilayer with the Pro1213 to Cys1236 segment constructed as an α-helix based on the results of the TM helix prediction algorithms (Fig. S1). The combined trajectory from three 500 ns simulation replicates revealed that the Cα atoms of residues Tyr1215 to Cys1236 form the 22-residue helix that remains embedded in the membrane throughout the simulation (Fig. 2A).

**Figure 2:**
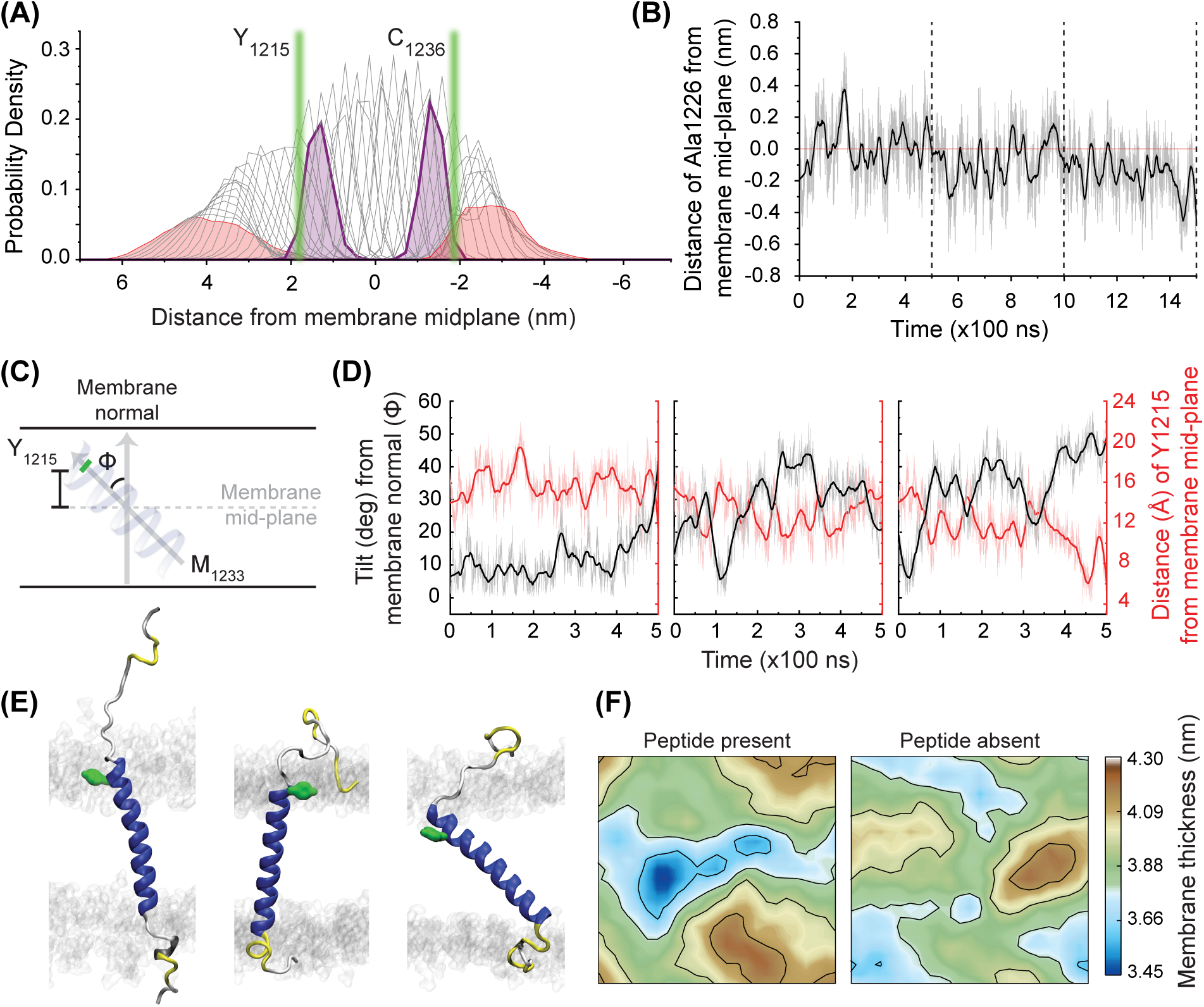
Determining membrane boundaries of the spike TM helix from atomistic simulations. **(A)** Probability density distributions (gray) of the Cα atoms for each of the 54 residues of the simulated polypeptide calculated from the 1.5 μs long combined simulation trajectory with the two terminal Ser residues (shaded red). Tyr1215 and Cys1236 (shaded purple) form the ends of the expanded consensus TM helix in Fig. 1B). The median of the centroids of the probability density distributions of phosphate from the lipid bilayer are shown as vertical green lines. It can be seen that the residues of the expanded consensus TM helix stay within the membrane. **(B)** Distance of the Cα atom of Ala1226 (the central residue of the expanded consensus helix) from the membrane midplane (red line) across the combined 1.5 μs trajectory (gray, 400 point average overlaid in black). The vertical dashed lines indicate ends of the three simulation replicates. **(C)** The tilt of the TM helix (angle between the membrane normal and the vector between the center of masses of the residues Met1233 and Tyr1215) and the distance of Tyr1215 from the membrane midplane should be anticorrelated for straight helices. **(D)** The TM helix tilt (black) and the distance of Tyr1215 from the midplane (red) from simulations shows such anticorrelation (simulation data as light trace and its 400 point average as the dark trace). **(E)** Snapshots of the three most populated conformations from each of the replicate simulations that were coarse-grained as T1, T2 and T3 (left to right). Side chain of Tyr1215 used in several simulation analyses is shown in green. The rest of the peptide backbone is colored according to secondary structure (helix in blue, turns in yellow and coil in white). In gray are the lipid headgroups. **(F)** The average membrane thickness in the simulation box (8 nm by 8 nm) when the TM helix is present (left panel) and without the TM helix (right panel). The last 25 ns data from a single replicate have been averaged. The peptide thins the membrane around it.

The membrane-embedded TM helix segment was found to be highly dynamic in the atomistic simulations. The Cα atom of Ala1226, which represents the approximate center of the TM helix moved more than 1 nm along the axis normal to the bilayer during the course of the combined trajectory centered at the bilayer (Fig. 2B). In addition to bobbing up and down, the TM helix tilted more than 50° with respect to the membrane normal and occasionally underwent bending (Fig. 2C-2E). Despite the occasional bending, the angle of tilt of the TM helix was negatively correlated with the distance of the Tyr1215 from the center of the bilayer (Fig. 2D). Moreover, the TM helix gained and lost secondary structure at both its N- and C-termini (Fig. S2). This means that the TRACS also did not remain helical across the simulations. The Cα atom of the Lys1211 snorkeling at the membrane boundary was pushed away from the lipid headgroups due to loss of helicity at the N-terminus of the spike TM helix (Fig. S3). Collectively, the dynamics of the TMD, especially the tilted conformations, created packing defects in the membrane, which caused local membrane thinning (Fig. 2F). The results from atomistic simulations justify the need for the separate nomenclature for TRACS, which is shorter than, but similar in sequence (*23*) to the <u>m</u>embrane <u>p</u>roximal <u>e</u>xternal region (MPER) from the HIV fusion protein, gp41 (*46*). However unlike the MPER that remains proximal to the membrane (*34, 46*), TRACS embeds in the bilayer during the simulations and so should not be considered completely analogous to MPER.

Although, Tyr1215 to Cys1236 stays membrane-embedded throughout the simulations and can be thought of as a minimal spike TM helix (Fig. 2A), the tilting and bobbing motions over tens of nanoseconds mean that both the sequence of the complete helical segment embedded in the membrane as well as its conformations are variable. A single representative conformation was then chosen from each replicate trajectory for further simulations (Methods). This procedure resulted in TM helices with the following sequences: Trp1212-Cys1236 (T1), Tyr1215-Cys1241 (T2), and Lys1211-Met1237 (T3) (Fig. 2E). We course-grained the three chosen TM segments using the Martini force field and tested their ability to associate into trimers, reflecting the stoichiometry of the SARS-CoV-2 spike protein (*3, 4*).

### TM helices assemble into symmetric and asymmetric trimers

The coarse-grained Martini force field has been widely used to investigate oligomerization in lipid membranes (*47–50*) because it permits greater sampling in a given time (*49*). Here, coarse-graining facilitates the study of homotypic interactions as they emerge among the spike TM helices. Additionally, the force field does not allow secondary structure to vary much over the course of the simulations (*51*), which we used to our advantage to examine the differences in association propensity of TM helices (T1, T2 and T3) formed by stretches of residues shifted in sequence (Fig. S4).

First, we modeled the 25 amino acid T1 sequence, Trp1212-Cys1236 (Fig. 3A), as an ideal α-helix, and examined its potential to self-assemble into trimers using Martini simulations. 100 replicates of unrestrained trimerization simulations, i.e., each with three T1 helices, were simulated for 4 μs each in a homogenous POPC membrane (Table S1) following a previously validated method (*48*). The last frame in 53% of the 100 simulation trajectories revealed a trimer, while the remaining 47% showed dimeric association (Fig. 3B). We analyzed the trimers by clustering the snapshots obtained by combining the last 50 ns from all trajectories, whose last frame showed a trimer. Using a 5 Å RMSD cutoff (Methods), 50 clusters were obtained (Fig. S5), which in itself is suggestive of a highly dynamic trimer population. Moreover, the diverse trimers are populated to a similar degree in the T1 simulations (Fig. S5) indicating marginal stability of these trimeric conformations, suggestive of an inherently dynamic system that might allow inter-convertibility between the conformations.

**Figure 3:**
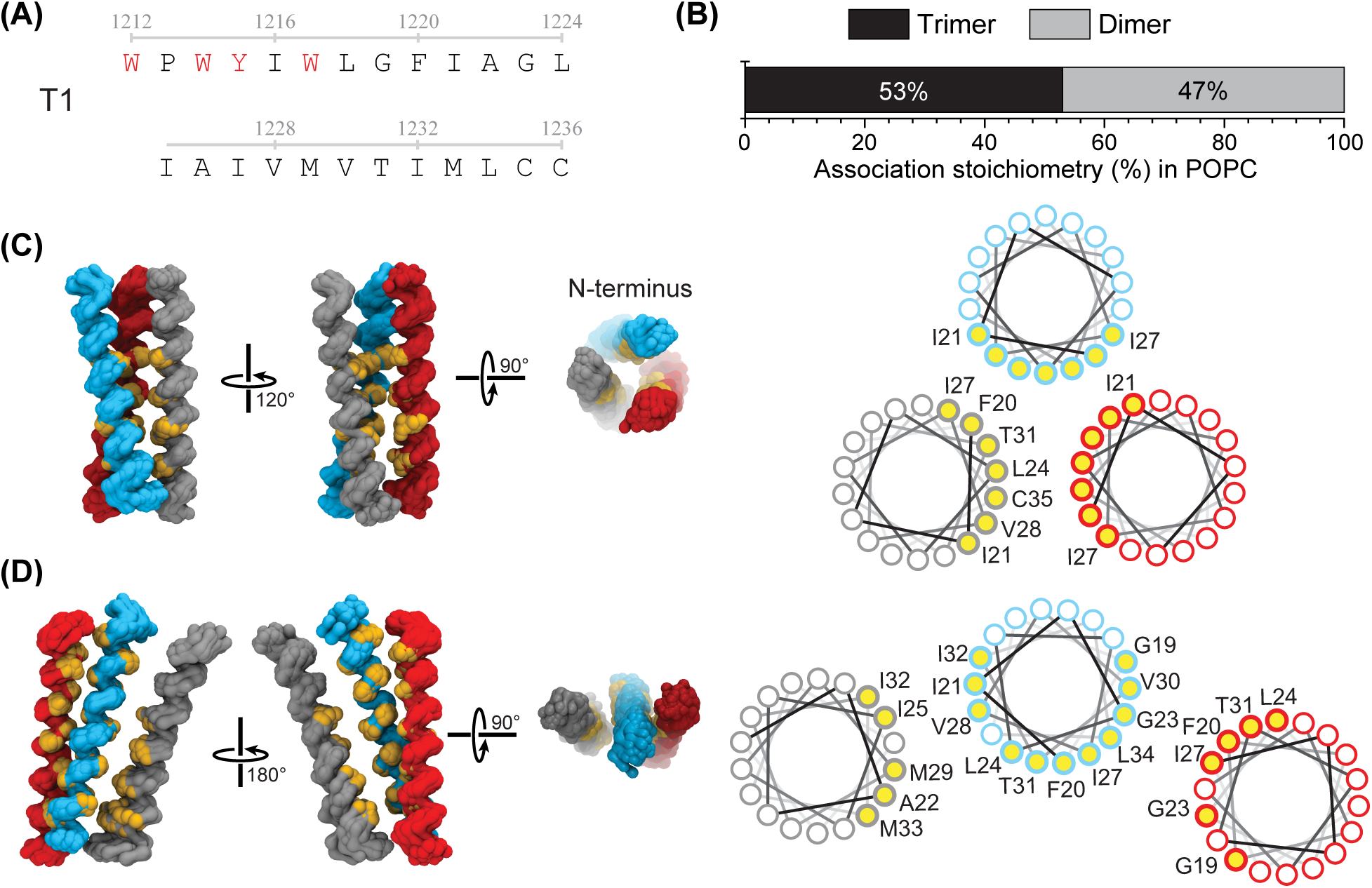
Spike TM helix forms both symmetric and asymmetric trimers. **(A)** Sequence of the T1 TM helix of 25 residues (aromatic residues comprising the N-terminal tryptophan-rich aromatic conserved stretch (TRACS) in red). **(B)** Stoichiometry of the self-associated T1 helices obtained from the last frame of 100 replicate trimerization simulations. T1 packing in the **(C)** symmetric and **(D)** asymmetric trimeric clusters. The three helices are shown in gray, cyan and red. The interacting residues are shown in yellow. The residues at the interface are shown and labelled in the helical wheel diagrams (same coloring as in the structures). For the symmetric interface, all the interacting residues are labelled only on the gray helix. Their identity can be inferred from the labelled end residues on the other two helices. Residue numbering follows the numbering in full length spike (see **(A)**) minus 1200 for brevity (e.g. G1219 is G19).

### Three overlapping motifs facilitate TM assembly

Two of the most populated trimeric conformational clusters were markedly different from each other. One of these clusters exhibited three-fold symmetry (Fig. 3C), while the other was asymmetric (Fig. 3D). The symmetric trimer exhibiting a left-handed supercoil was compact and was stabilized by homotypic interactions between the β-branched hydrophobic residues (F1220, Ile1221, Leu1224, Ile1227, Val1228) occupying one face of the helix (Figs. 3C and S6). In contrast, the asymmetric trimer was expanded, with two helices exhibiting dimerization using the GXXXG (Gly1219 to Gly1223) motif (*52–54*), otherwise known as the GAS_right_ motif (*55*) and the third helix docked onto this dimer, interacting over several hydrophobic residues (Figs. 3D and S7). Analysis of the contact maps revealed that the N-terminal conserved Trp residues that comprise the TRACS do not interact with each other and hence, do not participate in the homotypic association of T1 (Figs. S6 and S7).

The other highly populated clusters (Fig. S8) displayed a range of conformations that were intermediate between the compact symmetric trimer, and the expanded asymmetric trimer. With the exception of the symmetric trimer, which was compact at both the termini (Figs. 3C), all the clusters were observed to be more compact at the C-terminus than at the N-terminus (Figs. S8 and 3D). We found that a different GAS_right_ motif, formed by Ala1222 and Ala1226, stabilizes some of the intermediate conformations (Fig. S8). Therefore, some but not all the trimers from the T1 simulations matched the symmetry of the soluble ectomembrane domain of the spike.

### N-terminal TRACS is detrimental to the SARS-CoV-2 spike TMD trimer

We repeated the trimerization simulations with two more constructs derived from the all-atom simulation results, T2 (Tyr1215-Cys1241) and T3 (Lys1211-Met1237) (Fig. 4A), offset in sequence by four residues. Unlike the 25-residue long T1 construct, the T2 and T3 helices were comprised of 27 residues each, and only T2 lacked the complete TRACS (Fig. 4A). Coarse-grained T2 was modelled with ideal α-helical parameters like in the case of T1, however, T3 exhibited a slightly bent conformation and hence atomistic coordinates were used to generate a coarse-grained model for T3 (see Methods). 100 replicate Martini simulations were performed for both T2 and T3. As in the T1 simulations, each replicate 4 µs trajectory had three copies of a given construct (T2 or T3) in a homogenous POPC membrane (Table S1).

**Figure 4:**
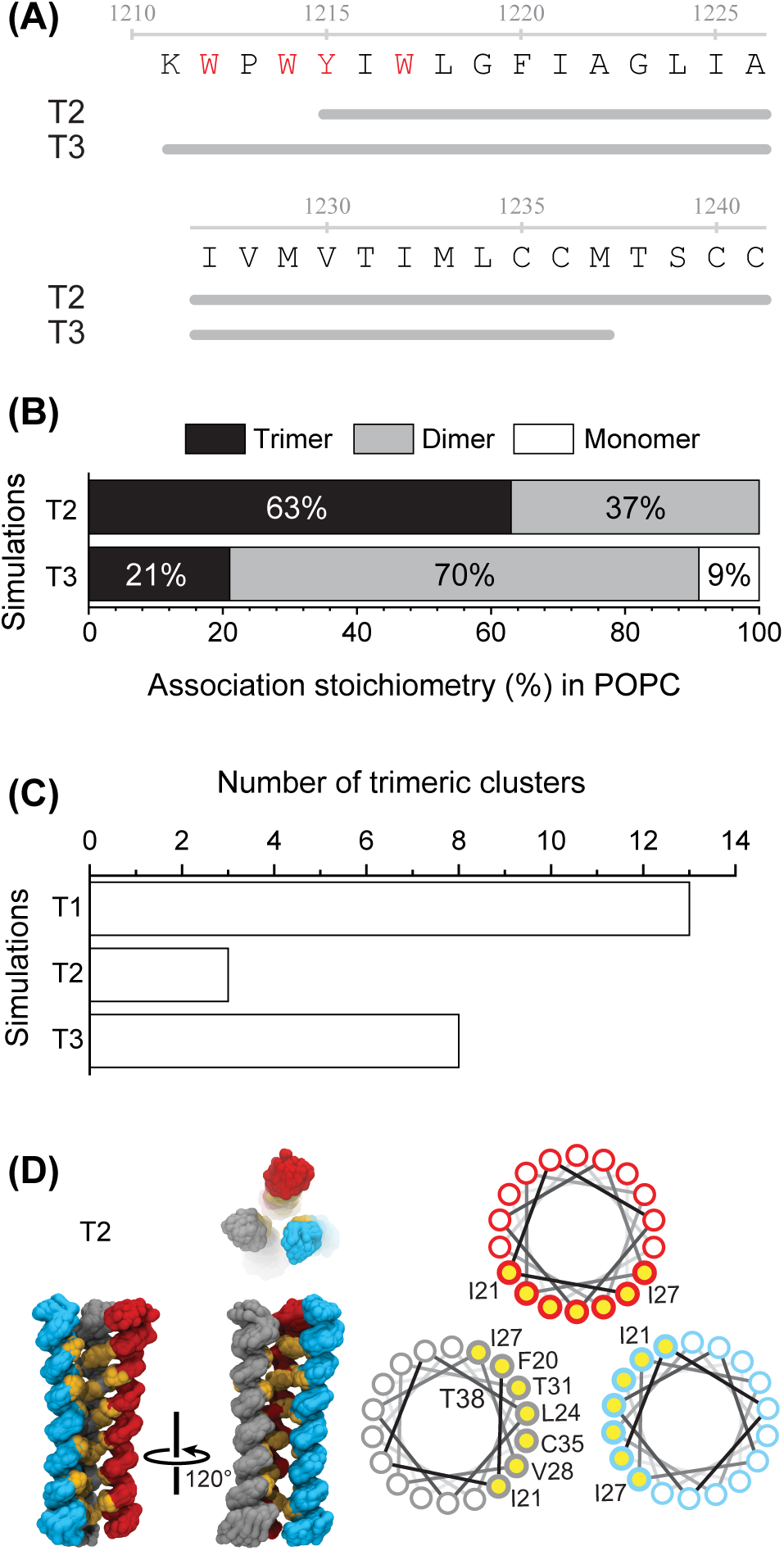
Sequence of the membrane-embedded spike TM helix affects self-assembly. **(A)** Gray lines below the spike sequence mark the 27 residues forming the T2 and T3 constructs. (TRACS in red). **(B)** Stoichiometry of the self-associated TM helices obtained from the last frame of 100 replicate trimerization simulations. **(C)** Number of the major trimeric clusters (see **Figs. S5 and S9**) of TMD conformations for the three simulated constructs. **(D)** The largest T2 cluster is similar to the symmetric cluster in Fig. 3C. Colors and labelling scheme same as in Fig. 3C. 63% of the final snapshots of the T2 replicates showed trimers (Fig. 4B). More importantly, T2 simulations displayed reduced structural diversity, with only three major structural ensembles observed after clustering the last 50 ns of all replicate trajectories that ended in trimers (Figs. 4C and S9A). The most populated cluster of the T2 simulations showed a symmetric trimer (Fig. 4D), with a nearly identical interaction surface as the T1 symmetric trimer (Fig. 3C), but extended by one helical turn at the C-terminus (Figs. 4D and S10). This extended interaction surface and the absence of two of the bulky Trp residues of the TRACS at the N-terminus of T2 resulted in improved packing of the helices (Fig. 4D) compared to T1 (Fig. 3C).

T3 showed the least trimerization (21%) among the three constructs and roughly 10% trajectories ended without a single T3-T3 association event (Fig. 4B). Also, trimeric conformations from all the trimerized simulations in T3 were diverse, similar to T1 (Figs. 4C and S9B). This structural diversity suggests that the inclusion of the bulky Trp from TRACS into the TM helix may increase TMD dynamics. It might be argued that differences in starting conformation (slightly bent and tilted helix in T3 as compared to T1 and T2), size of the simulation box, or peptide to lipid ratio contribute to the loss of trimerization propensity in T3. However, we systematically negated factors other than the inclusion of the TRACS sequence that could contribute to the decrease in trimerization propensity in T3. The most populated T3 trimer (Fig. S11) had straight helices despite the bent starting conformation (Fig. S4 and S12) due to the similarity between the default strength of the backbone dihedral restraints (K_BBBB_ = 400 kJ mol^-1^; (*51*)) and the strength of the peptide-lipid interactions in the Martini force field. Moreover, by definition the backbone dihedrals are weaker at the ends of the helix (*51*). So, slightly bent helices can straighten during the course of equilibration, as reported before in a different context (*56*). Another difference between the T1, T2, and T3 simulations was the size of the simulation box, which differed to accommodate a 6.5 nm inter-helix starting distance between the three copies of the TMD (Fig. S4). However, given our estimate of the diffusion constant of the helices (Fig. S13), helix collision frequency is unlikely to be a significant factor. While it is conceivable that a small variation in the lipid/protein ratio could be relevant, there is no evidence regarding such an effect. Trp residues seem to prefer to interact with the lipid headgroups and not participate in interhelical protein-protein interactions (Figs. S6 and S7). Therefore, the conservative interpretation of the results is that the sequence of the membrane-embedded portion of the peptides plays a big role in trimerization, and embedding the N-terminal TRACS in the membrane disfavors homotrimerization of the SARS-CoV-2 spike TMD.

### Cholesterol interaction discriminates between length of the TM helix in the membrane and utilizes the same residues as those involved in helical self-assembly

The N-terminus of the spike TM helix harbors a cholesterol (chol) binding sequence (KXXXYXXL) called CARC (*57*), which is an inverted chol recognition amino acid consensus (CRAC) motif (*58*). Also, both the SARS-CoV-2 viral envelope and the plasma membrane to which it fuses, have chol in them (*59, 60*). Hence, by repeating the Martini trimerization simulations in a POPC:chol membrane (Methods) tested the effect of the lipid environment on the association of spike TM helices. The same T1, T2, and T3 were used, and as before, three copies of each were simulated for 4 μs per trajectory (Table S1). 50 replicate trajectories were simulated. The trimerization propensity for T1 remain unaltered in the presence or absence of chol (Figs. 3B and 5A). However, T2 and T3 both displayed a 30% reduction in trimerization (Figs. 4B and 5A). Surprisingly, both T2 and T3 interacted with chol to a similar degree (Figs. 5B), despite T1 and T3 harboring the complete CARC motif.

**Figure 5:**
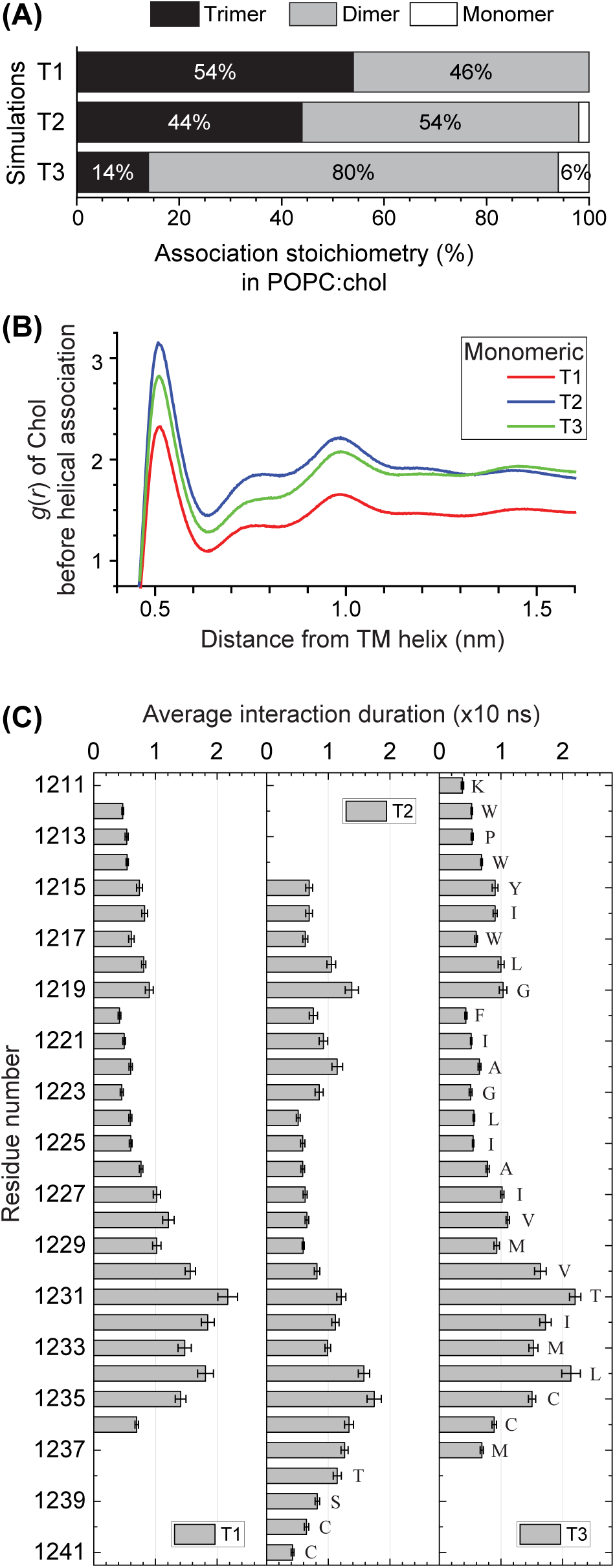
Cholesterol (chol) prefers to interact at the C-terminus of the spike TM helices. **(A)** Stoichiometry of the self-associated TM helices obtained from the last frame of the 50 replicate trimerization simulations in POPC:chol membranes. (Compare with **Figs. 3B** and **4B**). **(B)** The radial distribution function, *g(r)*, of cholesterol around monomeric helices (mean from 10 trajectories, see **Fig. S14** for the extended plot with error bars, where *g(r)* tends to 1 at longer distances). **(C)** Average interaction duration of cholesterol with each residue of the three TM constructs before the occurrence of any self-association event in the trajectory (mean ± SD averaged over 14 replicate simulations). The residues are labelled only on one of the three constructs.

Analysis of the residue-specific contacts of chol with monomeric TM helices revealed that the main site for chol interaction was not the predicted CARC motif at the N-terminus. Instead, chol interacted with the C-terminal half of the helix, and all the three sequences showed similar interaction with chol (Fig. 5C). This observation is consistent with a surface plasmon resonance (SPR) spectroscopy experiment of SARS-CoV-2 spike, which found minimal chol interaction at the predicted CARC motif (*61*). We did not find substantial differences between the diffusion of the T1, T2, or T3 helices (Fig. S15), the time to the first interaction (Fig. S16), or even the number and occupancy of interacting chol molecules per helix (Fig. S17). Additionally, the trimers formed by the spike TMD in the presence of chol are structurally similar to those formed in pure POPC bilayers (Figs. S18). Thus, the shorter T1 helix (25 residues) is less perturbed by chol and can trimerize as in POPC, unlike the longer, 27-residue T2 and T3. However, more work is required to completely understand the effects of chol on the degree of self-association of the spike TMD.

Nevertheless, we mapped the chol-bound conformations of the spike TM helix back to their atomistic coordinates (Methods). Both α and β faces of chol (*57*) make transient (∼15 ns) interactions with the TM helices in the simulations (Fig. 5C). Frequently observed sites of chol interaction at the C- and the N-termini are reported in Figs. S19 and S20, respectively. Despite the fact that the chol interacting region on the helix (Fig. 5C) overlaps with some of the residues involved in stabilizing the self-associated conformations (Fig. 4D), the final structures in the two lipidic environments are comparable (Figs. S18), highlighting the transient nature of the chol interaction. However, transient obstruction of the C-terminal half of the helix by chol (Figs. 5C and S17) prevents the C-terminus from initiating self-association, thereby influencing the initial events that occur upon the first encounter of TM helices. Unlike in pure POPC (Fig. 6A), the N-terminal residues generally initiate self-association in the presence of chol (Fig. 6B).

**Figure 6:**
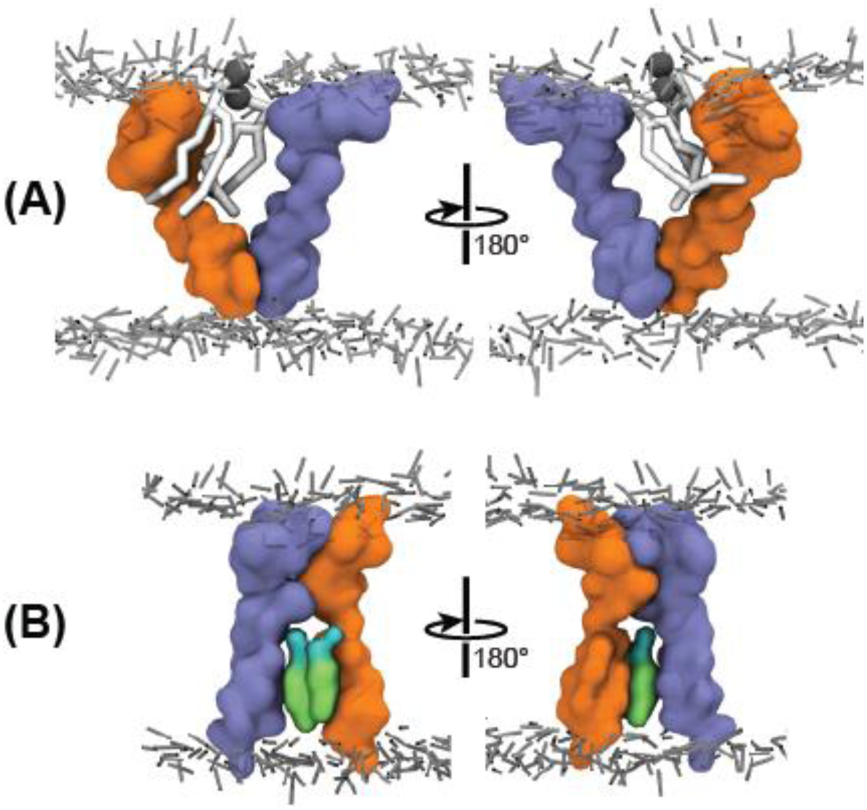
Chol alters the position of emergence of homotypic interactions. Representative snapshots of the first self-association event from trimerization simulations in **(A)** homogenous POPC membranes (C-terminal inter-helix interaction) and in **(B)** POPC:chol membranes. (N-terminal inter-helix interaction). Only the two interacting helices are shown in orange and violet, and, only the intervening POPC (white) and chol (green) molecules are shown. Lipid headgroups are shown as dark lines.

### Spike TM helix dimerization independently reveals mirrored super-helical handedness

The T3 simulations had an interesting trimeric ensemble, not seen in T1 or T2, which can be thought of as a right-handed dimer and a left-handed dimer sharing the central helix (Fig. S11). Moreover, 70% of all the T3 simulations ended dimeric in POPC membranes (Fig. 4B), increasing to 80% in chol-doped POPC membranes (Fig. 5A). The peculiar trimeric conformation and large dimeric fraction in T3 motivated us to investigate the dimerization of the TM helices in more detail. We simulated TM helices formed by two sequences offset by two residues, D1 (Trp1214-Cys1236) and D2 (Trp1212-Leu1234) that encompass the T3 sequence (Fig. 7A). These two 23-residue TM helices also include the minimal fragment that remained helical in the atomistic simulations (Fig. 2A). D1 and D2 were modelled as ideal α-helices and two copies of each were simulated using Martini in a POPC bilayer. 75 replicates were simulated, each for 1.5 μs (Table S1). The last frames of the replicates of the D1 and D2 constructs showed nearly 50% and 40% dimerization, respectively (Fig. 7B). However, clustering the conformations from the last 50 ns of the successfully dimerized trajectories, as done for the trimers, revealed strikingly different dimers (Figs. 7C-7D). The two dimeric ensembles were mirrored in their super-helical handedness, which is remarkable as D1 and D2 are offset by only two amino acids. D1 displayed classical right-handed dimers, stabilized by the GXXXG motif involving Gly1219 and Gly1223 (Fig. 7C). The D2 sequence formed a left-handed dimer, stabilized by an extensive interaction surface utilizing the LXXVIXXT segment (Fig. 7D). These results from dimerization simulations helped us dissect the two interaction surfaces seen in the spike TMD trimers, where one interface is formed by the GXXXG motif and the other is formed by the stretch of β-branched hydrophobic residues, LXXVIXXT that occupy one face of the helix (Fig. 7E). Reassuringly, the two interfaces identified in these dimerization simulations were comparable to those in the dimeric ensembles generated from the T3 simulations (Fig. S21) where 70% of the replicates ended in a dimer.

**Figure 7:**
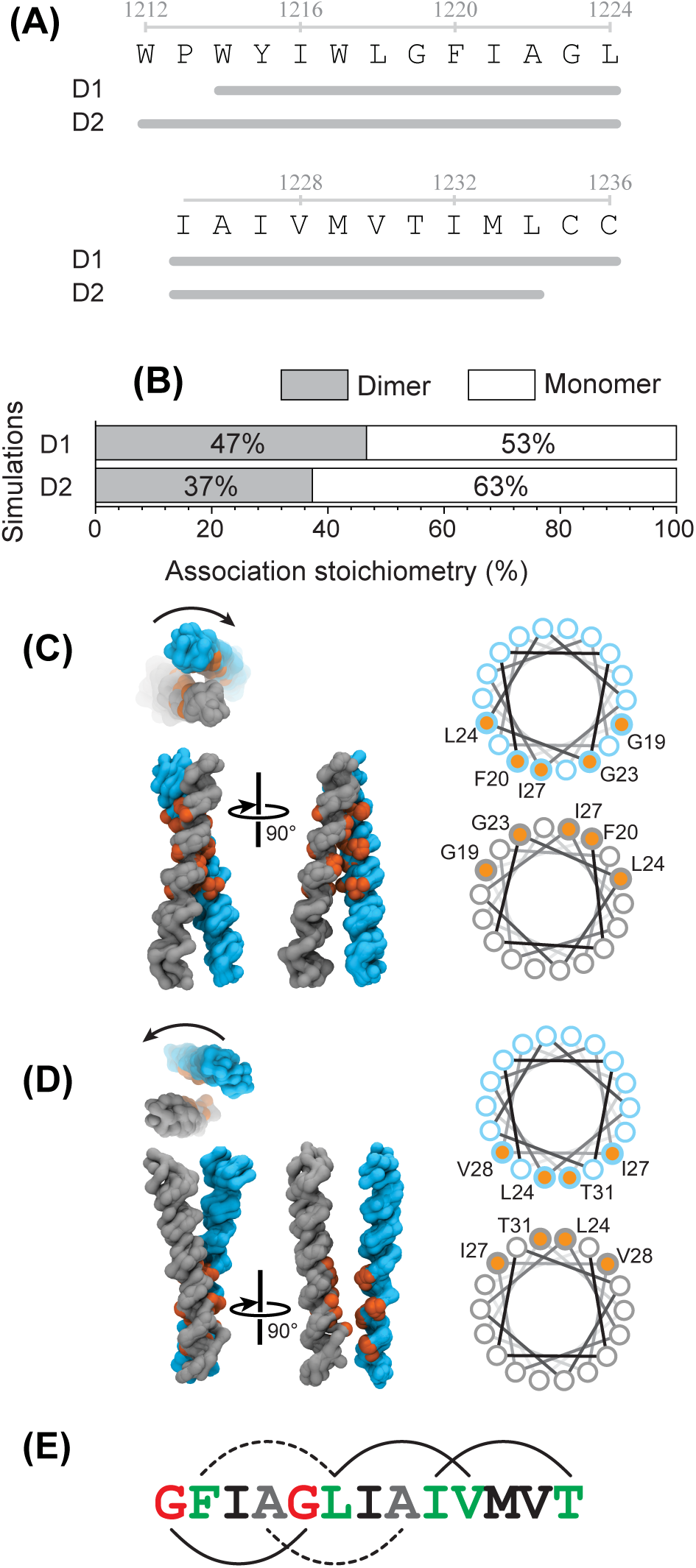
Dimerization simulations reveal two association interfaces that overlap to stabilize TM trimers. **(A)** Sequence of the D1 and D2 segments marked in gray lines. **(B)** Stoichiometry of the self-associated TM helices obtained from the last frame of the 75 replicate dimerization simulations. The major dimer conformation in the D1 simulations **(C)** shows right-handed super-helical dimerization and the D2 simulations **(D)** shows left-handed super-helical dimerization (indicated by curved arrows). **(C, D)** Helices are colored in cyan and gray on both the structural ensembles and the helical wheel diagrams. The interacting residues are shown in orange and also labelled on the helical wheel diagram. For brevity, residue numbering is that of the full length spike protein (see (**A**)) minus 1200. **(E)** The residues of the two identified dimerization motifs (GXXXG in red and LXXIVXXT in green) occupy i and i+4 residues and occur on the same face of the helix, as they do in the AXXXA motif (gray). Dotted lines mark the interfaces that are absent from the predominant dimer clusters, but seen in the trimeric conformations.

## DISCUSSION

### Dynamics of the spike protomer TM helix could destabilize the viral envelope and aid fusion

Non-consensus prediction of the spike protomer TM helix boundaries across different prediction servers agrees with the dynamic nature of the TM helix in atomistic simulations, indicating that the single TM helix may not fit a ‘conventional’ biological membrane. Specifically, the TM helix gained and lost helicity at both the N- and C-terminal membrane boundaries in a POPC bilayer, and the N-terminal TRACS (tryptophan-rich aromatic conserved stretch) was more dynamic and made more frequent excursions outside the bilayer. Loss of helicity correlated with the TM helix becoming perpendicular to the membrane, while gain of helicity allowed more residues to be buried in the membrane and a concomitant helical tilting of up to 55° (Fig. 2D). Such a high tilt is consistent with previous experiments on individual viral TM helices (*62*) and simulations of the functionally equivalent gp41 TMD from HIV (*37*). In the atomistic simulations, the TM helix dynamics induced a local disturbance in the lipid bilayer (Fig. 2F) due to constraints of hydrophobic matching (*63*). Destabilization of the viral envelope caused by such dynamics could decrease the kinetic barrier to fusion with the host membrane, as suggested for enveloped viruses (*64*).

The ectomembrane N- and the C-termini of the 54-residue simulated spike segment (containing the membrane-embedded TM helix) remained unstructured in atomistic simulations. This lack of secondary structure on both sides of the viral envelope is expected as the segment is not resolved in the cryo-ET studies of the SARS-CoV-2 spike (*65–67*). The proximity of the conserved Cys residues at the C-terminus of the simulated segment to the lipid headgroups (Fig. 2A) could promote its palmitoylation, which has been experimentally shown to disrupt the spike-mediated SARS-CoV-2 membrane fusion (*68*).

### Multiple overlapping and conflicting dimerization motifs determine the TMD conformation

The sequence motifs present in the spike TM helix interaction interfaces in the dimer simulations (Fig. 7E) were very similar to those seen in the trimer simulations. One of these interfaces is formed by the well-studied GXXXG or GAS_right_ motif (*52, 55, 69*). This motif is overrepresented in the naturally occurring TM helix sequence space (*70*) and promotes TM helix dimerization through a combination of hydrogen bonding and van der Waals packing interactions (*71, 72*). The other interface extends throughout one face of the TM helix and is comprised of β-branched residues (LXXIVXXT). Similar sequences have been previously shown to stabilize TM dimerization (*69, 73, 74*). The trimeric spike TMD is stabilized by these two overlapping dimerization interfaces and an interplay between these two sequence motifs dictates the final conformations observed. Additionally, a second GAS_right_ motif: AXXXA stabilizes intermediate structures of the trimeric TMD that are compact and closer to the three-fold symmetry at the C-terminus, and open and splayed at the N-terminus. Thus, the sequence of the spike TM region encodes multiple interaction motifs. Their relative contributions determine the conformation of the TMD, which is dependent upon whether they are accessible for interaction in the membrane-embedded TM-fragment. Another contributing factor is the location of these motifs with respect to the lipid bilayer, as the TM helix can move by 1 nm with respect to the bilayer (Fig. 2B).

### The role of the symmetric and asymmetric spike TMD trimers in fusion

Two different types of conformations were found for the trimeric spike TMD. First was a three-fold symmetric conformation (Fig. 4D) consistent with the symmetric ectomembrane structure in the resting pre-fusion spike trimer (*1, 4*). This conformation is stabilized by a LXXIVXXT interface, with minor variation occurring in the T1 and T2 constructs (Figs. 3C and 4D). This robustness of the symmetric TMD homotrimer may contribute to the stabilization of the pre-fusion trimer of the spike protein and keep it quiescent until the time when S2 begins reorganizing towards the post-fusion conformation (Fig. 1A).

Asymmetric TMD is the other significantly populated trimeric conformation in the T1 (Fig. 3D) and T3 (Fig. S11) simulations. Fusion proteins from enveloped viruses, such as the SARS-CoV-2 spike protein, must transit through an intermediate state between the symmetric pre- and post-fusion conformations where the symmetry is lost (Fig. 1A) (*75, 76*). This loss of symmetry has been suggested to occur in the juxtamembrane part of the spike protein as exemplified by the prehairpin intermediate (Fig. 1A). Since the TMD is connected to the juxtamembrane segment, the asymmetric conformations of the TMD seen in simulations, may locally stabilize and facilitate the formation of such an intermediate. Combining these results from the simulations, we next postulate a model of pre- to post-fusion conformational transition of the spike TMD (Fig. 1A).

### Working model of the dynamics of SARS-CoV-2 spike TMD during fusion

The symmetric trimer observed in T1 simulations (Fig. 3C), that becomes longer and more compact in T2 (Fig. 4D) when the TRACS in not membrane-embedded, corresponds to the pre-fusion conformations of the spike TMD. The large pre- to post-fusion conformational transition of the full spike protein has intermediate conformations (Fig. 1A) triggered by spike-host protein interactions, which could force the TRACS into the envelope. A TM helix with its TRACS inserted into the viral envelope is akin to the T3 construct. T3 simulations show that the incorporation of TRACS into the TM helix reduces its propensity to trimerize (Fig. 4B). Further, the trimeric conformations that are observed in T3 (Fig. S11) are asymmetric like those seen in T1 (Fig. 3D). Such asymmetric conformations are likely to facilitate a transition to the experimentally observed asymmetric prehairpin intermediates (*75*) (Fig. 1A). Asymmetric TMD trimers may dissociate via dimeric conformations seen in the D1 and D2 simulations (Fig. 7) driving the spike towards its post-fusion structure. Finally, local perturbations to the membrane thickness caused by the bobbing and tilting motions of the monomeric TM helix as seen in the atomistic simulations (Fig. 2) should help the viral envelope fuse with the host membrane. Overall, the many conformations observed in our simulations indicate that the TMD dynamics would complement and aid the large conformational change occurring in the ectomembrane domains of the spike protein during fusion.

### The effects of cholesterol on viral fusion

The lipid composition of the virus envelope depends on the host membrane from which it originates (*59*). Coronaviruses, such as SARS-CoV-2, assemble at the ERGIC (endoplasmic reticulum-golgi intermediate compartment) (*77, 78*), so their envelopes are expected to be similar in composition to the ERGIC membrane comprising ∼15 mol% chol (*60*). Mature SARS-CoV-2 fuses with plasma membrane (PM) upon infection and that could have up to 40 mol% chol (*60*). Thus, we chose an intermediate chol content of 30 mol% in the POPC:chol Martini simulations to study the influence of chol on the interacting TM helices. The fraction of the trimerization observed, but not the TMD conformations, was affected upon addition of chol to the membrane. The C-terminal half, and not the CRAC motif, of the SARS-CoV-2 spike TM helix showed more affinity towards chol (Fig. 5C). Additionally, the position of initiation of the TM protein-protein interactions was affected because of an overlap between the chol-interacting residues and the residues that form inter-TM helix contacts (Fig. 6). In the context of fusion, where the lipids from the viral envelope mix with the host membrane, chol from the PM is expected to diffuse into the viral envelope as seen in simulations of HIV membrane fusion (*79*). Our simulations indicate that an increase in chol in the TMD environment (due to an inflow from the PM) could further weaken the association of the TM helices, which would drive the spike TMD towards its post-fusion conformation (Fig. 1A) where the TM helices do not exhibit homotypic interactions.

### Influencing the stability of the pre-fusion spike conformation by modulating TMD dynamics

Large changes to the trimerization of the spike TM helices were observed depending upon the presence and depth of TRACS in the membrane (Fig. 4B). The T2 helix that starts from Tyr1215 does not contain the complete TRACS (Trp1212-Trp1217). Analysis of the interaction surface in T2 reveals the Trp1217 residue does not participate in trimerization, although the Tyr1215 residue does form infrequent homotypic contacts by virtue of being on the same face of the helix as the extended FXXXLXXIVXXT trimeric interface. The T3 helices contain the complete membrane-embedded TRACS, and remain largely dimeric exhibiting negligible symmetric trimers. Consequently, membrane-embedding the helical TRACS may destabilize the trimeric spike TMD and in turn, the entire pre-fusion conformation of the trimeric spike protein.

The (de)stabilization of specific conformations of the spike in order to block its membrane fusion activity has been explored before. Small polypeptides and lipopeptides have been reported to function in a similar manner (*17–19*). Additionally, TRACS is an attractive target because it is conserved across coronaviruses (*20–22, 24*) and a molecule targeting the TRACS could be broad spectrum in its antiviral activity. We note that the sequence of TRACS is nearly identical to that of indolicidin (Fig. 8), which is an established antimicrobial peptide (*80, 81*) with demonstrated anti-viral activity (*82*). Since indolicidin is known to favorably insert into membrane-water interfaces (*83*), which is the location of the TRACS, we hypothesize that indolicidin may specifically target TRACS, disrupt the TMD assembly, and consequently that of the pre-fusion conformation of the spike protein. Thus, indolicidin is a potential molecular scaffold for early therapeutic intervention in SARS-CoV-2 infection.

**Figure 8:**
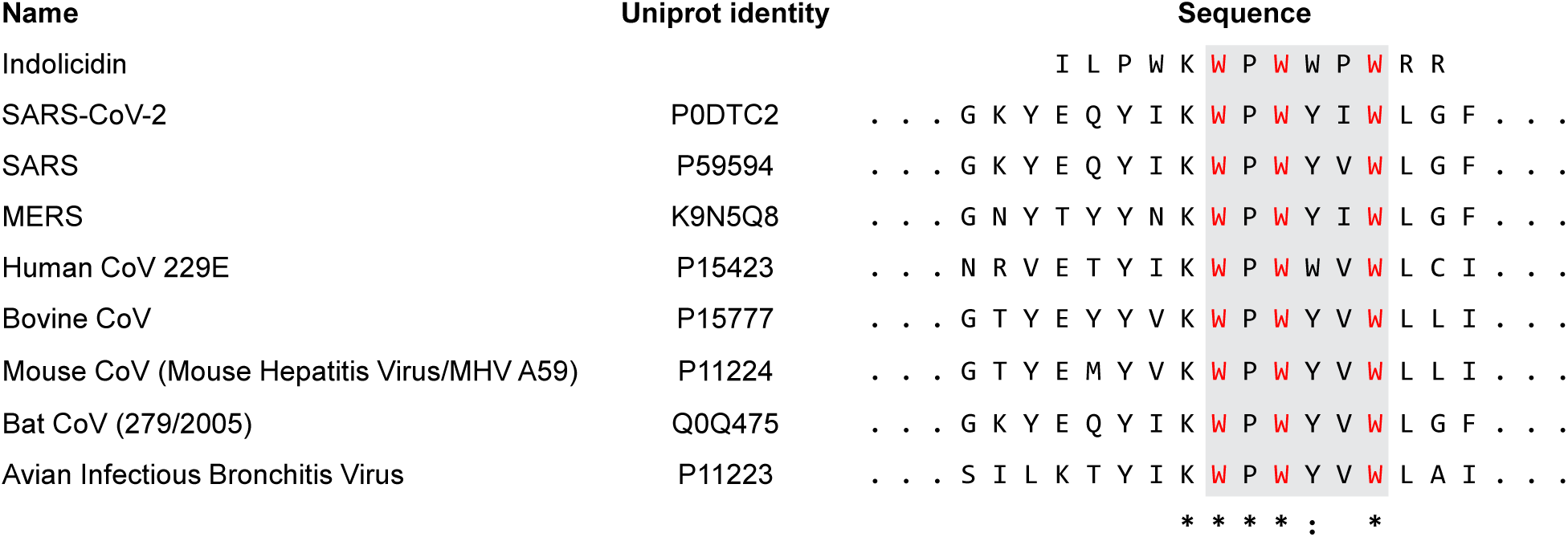
The TRACS and indolicidin sequences are similar. Manual sequence alignment of indolicidin with the TRACS (gray box) from a range of coronaviruses. Invariant Trp residues from TRACS are colored red, asterisks indicate complete identity and colon indicates partial conservation.

## CONCLUSIONS

The SARS-CoV-2 spike protein undergoes a large conformational change in order to enable the fusion of the viral envelope to the host cell membrane. Three helices, each contributed by one protomer of the homotrimeric spike, and together called the transmembrane domain (TMD), anchor the spike protein to the viral envelope. It is assumed that the TMD is a passive anchor that doesn’t actively participate in the spike conformational changes that enables viral fusion. We performed multiscale MD simulations of the spike TM helix in order to understand the populated structural ensembles and self-assembly dynamics of the TMD. Our atomistic simulations showed that a segment of the spike protomer embedded in the membrane and containing the putative TM helix is highly dynamic. The TM helix could melt and reform near the membrane boundaries and also bobbed and tilted in the membrane. We then coarse-grained the TM helical segments of three distinct conformations obtained from these atomistic simulations using Martini and performed trimerization simulations of each conformation in pure POPC and POPC:chol membranes. These simulations showed that there were multiple overlapping and conflicting interaction interfaces within the TM helices. The assembled TMD conformation depended on which of these interfaces participated in inter-helix interactions. A structural analysis of these conformations indicates that the TMD is not just a passive anchor but its dynamics may, in fact, facilitate the large conformational transitions occurring in the spike ectomembrane domains which are required for viral fusion. Among the highly populated conformations, a symmetric conformation mirrors the symmetry of the pre-fusion state of spike and is likely the resting pre-fusion conformation of the TMD. Another major conformation, which is asymmetric, may enable the TMD disassembly which is required to form the asymmetric spike prehairpin fusion intermediate. The presence of chol modulates the amount of trimerization but does not affect the types of conformations populated. In summary, the simulations show that dynamics are encoded in the spike TM helix sequence. The protomer dynamics enable the population of structurally distinct dimer and trimer conformations. Finally, the observed TMD conformations and dynamics are likely to promote the large conformational change occurring in the ectomembrane domains of the spike protein. Thus, molecules that alter spike TMD dynamics may reduce SARS-CoV-2 fusion and we suggest a molecular scaffold for such a therapeutic intervention.

## METHODS

### Generating the starting atomistic conformation

There are no structural coordinates for the pre-fusion spike transmembrane (TM) domain (TMD) in the protein data bank (PDB). Nor do we know the start and the end residues of the TM helix of a spike protomer. Ser1196 is at least 15 residues before the start of the hydrophobic stretch in the spike sequence. And on the other side of the viral envelope, the residues downstream of Ser1249 are known to comprise the C-terminal tail that exists inside the virus. Therefore, Ser1196-Ser1249, a 54 residue polypeptide containing the TM helix was simulated.

To assign secondary structure to the Ser1196-Ser1249 polypeptide, various secondary structure and TM helix prediction tools were used (Fig. S1A). The molefacture plugin in VMD (Visual Molecular Dynamics) (*84*) was used to model Pro1213-Cys1236 as the starting TM helix with ideal α-helical backbone torsion angles (*φ* = -57°, *ψ* = -47°), while rest of the polypeptide was modeled with *φ* = -60°, *ψ* = 30°.

The further processing of the polypeptide and preparation of the simulation box was done using the membrane builder of CHARMM-GUI (*85–87*). Since a segment of the spike was being simulated, its termini were charge neutralized by amidation at the C-terminus and by using a neutral amino group at the N-terminus. This polypeptide was then embedded in a planar, fully hydrated POPC (1-palmitoyl-2-oleoyl-glycerophosphatidylcholine) bilayer. This starting conformation comprising one copy of the 54 residue polypeptide, 203 lipid molecules, 22,966 water molecules (TIP3P) and 0.15 mM KCl was assembled in a cuboidal box of 8 nm x 8 nm x 14.5 nm that allowed the extended ectomembrane segment to be accommodated. The resulting initial configuration comprised 97,083 atoms (Fig. S1B).

All the simulations were performed using periodic boundary conditions in Gromacs 4.6.7 ((*88*), www.gromacs.org).

### Atomistic simulation of the single TM helix containing polypeptide

The equilibration and the production run of the simulation box were performed using the CHARMM36 force field (*89*) according to parameters provided in Gromacs compatible output files from CHARMM-GUI membrane builder (*85, 86*). The recommended six-step equilibration procedure (*86*) was followed. The first equilibration used 5000 steps of steepest descent energy minimization with position restraints on heavy atoms. This was followed by two NVT equilibration simulations and three NPT simulations. This stepwise equilibration provided an opportunity to relax the forces restraining the heavy atoms with every equilibration. The bonds involving hydrogen atoms were constrained with LINCS (*90*). Particle mesh Ewald (PME) summation (*91*) was used for the treatment of long range electrostatics, and the van der Waals interactions were cut off at 1.2 nm. Temperature was maintained at 303.15 K by coupling the system to the Berendsen’s thermostat (*92*) using a time constant (*τ*_t_) of 1 ps. Semi-isotropic pressure on the system was controlled at 1 bar with the Berendsen barostat (*92*) set at a coupling constant (*τ*_p_) of 5 ps. The time step was 1 fs throughout the equilibration, except for the last NPT simulation, where it was set at 2 fs.

The only difference between the parameters of the last NPT equilibration and the production run was the replacement of the Berendsen’s thermostat and barostat with the Nose-Hoover thermostat (*93, 94*) (303.15 K, *τ*_t_ = 1 ps) and the Parinello-Rahman barostat (*95*) (semi-isotropic presssure, 1 bar; *τ*_p_ = 5 ps). Three replicate production trajectories starting with different initial velocities generated using the *gen_vel* parameter in the mdp file and a unique seed for each of the three, were simulated for 500 ns each. Control simulations of identically sized hydrated POPC membrane were performed without the embedded polypeptide in three replicates of 100 ns each. A summary of all the simulations is provided in Table S1.

### Analysis of atomistic simulations

The simulations were visualized with VMD (*84*). The calculation of secondary structure, atomic distance, angle, density and RMSD were done using inbuilt Gromacs routines, *do_dssp* (Fig. S2), *dist* (Figs. 2B-2D), *angle* (Fig. 2D), *dens* (Figs. 2A and S3) and *rms*, respectively. The bilayer was centered before analyzing any of the motions of the peptide in the membrane plane to prevent artifacts arising from the translation motion of the bilayer with respect to the simulation box. GridMAT (*96*) was used to calculate the bilayer thickness (Fig. 2F).

Three structures (T1, T2, and T3), one from each replicate, were chosen from the atomistic simulations to use in coarse-grained Martini self-assembly simulations. To choose these structures, *do_dssp* derived assessment of secondary structure (Fig. S2) was manually analyzed for each replicate trajectory and several structures with long helical membrane-embedded conformations were identified from across the trajectory. Each of these conformations was used as a reference to calculate RMSD across the entire trajectory. The conformation that showed the least RMSD across the trajectory was chosen as being the representative of that trajectory.

### Coarse-grained self-association simulations

Coarse-grained self-assembly simulations were performed using the Martini 2.2 force field (*50, 51*). Three sets of trimerization simulations (T1, T2 and T3) each with three copies of identical helices were setup (details in Table S1). Since T1 and T2 were straight helices, they were built as idealized α-helices, while the bent and tilted T3 was built from its atomistic conformation. All three were coarse-grained to their Martini description using the *martinize* script (*50*). The DAFT protocol (*48*) was utilized to generate simulation systems where three in-plane rotated helices were placed equidistant from each other and their periodic images with a starting inter-helix center of mass (COM) distance of 6.5 nm. These were then embedded in a homogenous, planar POPC bilayer and solvated in Martini water (*97*). We used this procedure to generate 100 initial structures each for T1, T2 and T3 simulations. For the self-assembly simulations in cholesterol (chol) containing POPC membrane, the same starting structures prepared for T1, T2, and T3 were embedded in a POPC bilayer doped with 30 mol% chol and solvated in Martini water. No additional salt molecules were added. The dimerization simulations were performed by generating ideal α helices D1 and D2 from the TM sequence. As in the trimerization simulations, two copies of either D1 or D2 were placed 6.5 nm away from each other and their periodic images. The helices were embedded in a POPC membrane and solvated in Martini water. 75 starting systems were generated for each of the D1 and D2 (Table S1).

All the simulations were performed as described (*48, 50*). For equilibration, the starting conformations were energy minimized with 500 steps of steepest descent, followed by 100 ps of NPT with a 20 fs time step to bring temperature and pressures to their set value. Temperature was maintained by the *v*-rescale thermostat (310 K, *τ*_t_ = 1 ps) (*98*). Pressure was maintained using the Berendsen barostat (semi-isotropic, 1 bar; *τ*_p_ = 10 ps) (*92*). Coulombic interactions were cut-off beyond 1.2 nm, and the van der Waals interactions were switched to zero between 0.9 and 1.2 nm.

The production runs had the same simulation parameters, except for *τ*_p_, which was changed to 3 ps. Each trimerization replicate was simulated for 4 µs and each dimerization replicate was simulated for 1.5 µs. Details of the number of simulations for each ensemble and their duration are provided in Table S1.

### Analysis of the coarse-grained simulations

VMD (*84*) was used for visualizing trajectories and rendering graphics. Self-association stoichiometry of the TM helices was determined by the last frame of each trajectory. Helices were defined to be self-associated if the COM distance, as measured using Gromacs routine *dist*, was less than 1.2 nm (Figs. 3B, 4B, 5A, 7B). The radial distribution function (RDF), angle and mean square displacement were calculated by the Gromacs functions *rdf* (Figs. 5B and S14), *angle* (Fig. S12) and *msd* (Figs. S13 and S15). A linear fit of the MSD of the TM helices vs time plot was used to determine the diffusion coefficient for the TM helices (Figs. S13 and S15). To obtain the dimeric (in the dimerization simulations) and trimeric (in the trimerization simulations) structural ensembles, the last 50 ns of all the successfully dimerized and trimerized trajectories were extracted. These conformations were clustered with a one-to-one RMSD comparison across the self-assembled conformations using *g_cluster* with the -gromos flag (Figs. S5 and S9). Average contact maps were generated for each self-assembled conformation cluster using a 5 Å cutoff (Figs. S6, S7 and S10). ‘Interacting residues’ (marked in the figures in yellow) remained in contact in all the conformations in a cluster, i.e., had a value of 1 in the average contact map.

RDF of chol around individual TM helices was calculated before the first self-association event occurred, i.e., when the TM helices were still monomeric in the simulations. The final chol RDF value was an average determined from helices across 10 different randomly chosen trajectories, for T1, T2 and T3. PyLipID (*99*) was used for calculating chol-TM helix interactions (Figs. 5C and S17). To examine the chol binding modes in atomistic detail (Figs. S19 and S20), we *backmapped* (*100*) the chol-bound conformations of the TM helix extracted from PyLipID using the protocol recommended in the original report (*100*).

## Acknowledgements

This work was funded by the Department of Atomic Energy, Government of India through the Tata Institute of Fundamental Research from Project Identification No. RTI 4006. We acknowledge the NCBS high performance computing facility for computing resources. SG was supported in part by the Simons Foundation (Grant No. 287975), a grant from the National Supercomputing Mission (NSM) through the grant MeitY/R\&D/HPC/2(1)/2014 and SERB Grant: CRG/2021/004754. PB acknowledges the financial support in the form of DST-YOS Chair Professorship. SL was supported with a graduate fellowship from NCBS, TIFR and a bridging postdoctoral fellowship through the Simons Foundation (Grant No. 287975).

## Data availability

The starting coordinates for the three atomistic simulations and the coordinates for the three coarse-grained helices (T1, T2 and T3) are provided in SI.

## REFERENCES

1. C. B. Jackson, M. Farzan, B. Chen, H. Choe, Mechanisms of SARS-CoV-2 entry into cells. Nature reviews. Molecular cell biology 23, 3–20 (2022).

2. R. Peng, L. A. Wu, Q. Wang, J. Qi, G. F. Gao, Cell entry by SARS-CoV-2. Trends in biochemical sciences 46, 848–860 (2021).

3. F. Li, Structure, Function, and Evolution of Coronavirus Spike Proteins. Annu Rev Virol 3, 237–261 (2016).

4. J. Zhang, T. Xiao, Y. Cai, B. Chen, Structure of SARS-CoV-2 spike protein. Curr Opin Virol 50, 173–182 (2021).

5. A. C. Walls et al., Structure, Function, and Antigenicity of the SARS-CoV-2 Spike Glycoprotein. Cell 181, 281–292 e286 (2020).

6. Y. Cai et al., Distinct conformational states of SARS-CoV-2 spike protein. Science 369, 1586–1592 (2020).

7. D. Wrapp et al., Cryo-EM structure of the 2019-nCoV spike in the prefusion conformation. Science 367, 1260–1263 (2020).

8. D. J. Benton et al., Receptor binding and priming of the spike protein of SARS-CoV-2 for membrane fusion. Nature 588, 327–330 (2020).

9. A. C. Walls et al., Cryo-electron microscopy structure of a coronavirus spike glycoprotein trimer. Nature 531, 114–117 (2016).

10. R. N. Kirchdoerfer et al., Pre-fusion structure of a human coronavirus spike protein. Nature 531, 118–121 (2016).

11. L. Tai et al., Nanometer-resolution in situ structure of the SARS-CoV-2 postfusion spike protein. Proceedings of the National Academy of Sciences of the United States of America 118, (2021).

12. X. Fan, D. Cao, L. Kong, X. Zhang, Cryo-EM analysis of the post-fusion structure of the SARS-CoV spike glycoprotein. Nature communications 11, 3618 (2020).

13. A. C. Walls et al., Tectonic conformational changes of a coronavirus spike glycoprotein promote membrane fusion. Proceedings of the National Academy of Sciences of the United States of America 114, 11157–11162 (2017).

14. J. Pallesen et al., Immunogenicity and structures of a rationally designed prefusion MERS-CoV spike antigen. Proceedings of the National Academy of Sciences of the United States of America 114, E7348–E7357 (2017).

15. C. Wang et al., Antigenic structure of the human coronavirus OC43 spike reveals exposed and occluded neutralizing epitopes. Nature communications 13, 2921 (2022).

16. X. Song et al., Cryo-EM analysis of the HCoV-229E spike glycoprotein reveals dynamic prefusion conformational changes. Nature communications 12, 141 (2021).

17. R. D. de Vries et al., Intranasal fusion inhibitory lipopeptide prevents direct-contact SARS-CoV-2 transmission in ferrets. Science 371, 1379–1382 (2021).

18. S. Xia et al., Inhibition of SARS-CoV-2 (previously 2019-nCoV) infection by a highly potent pan-coronavirus fusion inhibitor targeting its spike protein that harbors a high capacity to mediate membrane fusion. Cell research 30, 343–355 (2020).

19. S. Xia et al., A pan-coronavirus fusion inhibitor targeting the HR1 domain of human coronavirus spike. Sci Adv 5, eaav4580 (2019).

20. Y. Liao, S. M. Zhang, T. L. Neo, J. P. Tam, Tryptophan-dependent membrane interaction and heteromerization with the internal fusion peptide by the membrane proximal external region of SARS-CoV spike protein. Biochemistry 54, 1819–1830 (2015).

21. J. Corver, R. Broer, P. van Kasteren, W. Spaan, Mutagenesis of the transmembrane domain of the SARS coronavirus spike glycoprotein: refinement of the requirements for SARS coronavirus cell entry. Virol J 6, 230 (2009).

22. Y. Lu, T. L. Neo, D. X. Liu, J. P. Tam, Importance of SARS-CoV spike protein Trp-rich region in viral infectivity. Biochem Biophys Res Commun 371, 356–360 (2008).

23. M. W. Howard et al., Aromatic amino acids in the juxtamembrane domain of severe acute respiratory syndrome coronavirus spike glycoprotein are important for receptor-dependent virus entry and cell-cell fusion. J Virol 82, 2883–2894 (2008).

24. E. Arbely, Z. Granot, I. Kass, J. Orly, I. T. Arkin, A trimerizing GxxxG motif is uniquely inserted in the severe acute respiratory syndrome (SARS) coronavirus spike protein transmembrane domain. Biochemistry 45, 11349–11356 (2006).

25. A. Walls et al., Crucial steps in the structure determination of a coronavirus spike glycoprotein using cryo-electron microscopy. Protein Sci 26, 113–121 (2017).

26. N. Mittal et al., Enhanced protective efficacy of a thermostable RBD-S2 vaccine formulation against SARS-CoV-2 and its variants. NPJ Vaccines 8, 161 (2023).

27. A. C. Walls et al., Unexpected Receptor Functional Mimicry Elucidates Activation of Coronavirus Fusion. Cell 176, 1026–1039 e1015 (2019).

28. Q. Fu, J. J. Chou, A Trimeric Hydrophobic Zipper Mediates the Intramembrane Assembly of SARS-CoV-2 Spike. J Am Chem Soc 143, 8543–8546 (2021).

29. W. Shi et al., Cryo-EM structure of SARS-CoV-2 postfusion spike in membrane. Nature 619, 403–409 (2023).

30. H. Yao, M. W. Lee, A. J. Waring, G. C. Wong, M. Hong, Viral fusion protein transmembrane domain adopts beta-strand structure to facilitate membrane topological changes for virus-cell fusion. Proceedings of the National Academy of Sciences of the United States of America 112, 10926–10931 (2015).

31. S. M. Costello et al., The SARS-CoV-2 spike reversibly samples an open-trimer conformation exposing novel epitopes. Nature structural & molecular biology 29, 229–238 (2022).

32. T. Mori et al., Elucidation of interactions regulating conformational stability and dynamics of SARS-CoV-2 S-protein. Biophysical journal 120, 1060–1071 (2021).

33. E. Dodero-Rojas, J. N. Onuchic, P. C. Whitford, Sterically confined rearrangements of SARS-CoV-2 Spike protein control cell invasion. Elife 10, (2021).

34. A. Piai et al., Structural basis of transmembrane coupling of the HIV-1 envelope glycoprotein. Nature communications 11, 2317 (2020).

35. J. Dev et al., Structural basis for membrane anchoring of HIV-1 envelope spike. Science 353, 172–175 (2016).

36. D. J. Benton et al., Influenza hemagglutinin membrane anchor. Proceedings of the National Academy of Sciences of the United States of America 115, 10112–10117 (2018).

37. L. R. t. Hollingsworth, J. A. Lemkul, D. R. Bevan, A. M. Brown, HIV-1 Env gp41 Transmembrane Domain Dynamics Are Modulated by Lipid, Water, and Ion Interactions. Biophysical journal 115, 84–94 (2018).

38. V. K. Gangupomu, C. F. Abrams, All-atom models of the membrane-spanning domain of HIV-1 gp41 from metadynamics. Biophysical journal 99, 3438–3444 (2010).

39. J. H. Kim, T. L. Hartley, A. R. Curran, D. M. Engelman, Molecular dynamics studies of the transmembrane domain of gp41 from HIV-1. Biochimica et biophysica acta 1788, 1804–1812 (2009).

40. A. Yu et al., A multiscale coarse-grained model of the SARS-CoV-2 virion. Biophysical journal 120, 1097–1104 (2021).

41. H. Woo et al., Developing a Fully Glycosylated Full-Length SARS-CoV-2 Spike Protein Model in a Viral Membrane. J Phys Chem B 124, 7128–7137 (2020).

42. L. Casalino et al., Beyond Shielding: The Roles of Glycans in the SARS-CoV-2 Spike Protein. ACS Cent Sci 6, 1722–1734 (2020).

43. C. T. Barrett, R. E. Dutch, Viral Membrane Fusion and the Transmembrane Domain. Viruses 12, (2020).

44. G. von Heijne, The membrane protein universe: what’s out there and why bother? J Intern Med 261, 543–557 (2007).

45. S. H. White, W. C. Wimley, Membrane protein folding and stability: physical principles. Annual review of biophysics and biomolecular structure 28, 319–365 (1999).

46. Y. Wang et al., Topological analysis of the gp41 MPER on lipid bilayers relevant to the metastable HIV-1 envelope prefusion state. Proceedings of the National Academy of Sciences of the United States of America 116, 22556–22566 (2019).

47. S. J. Marrink et al., Computational Modeling of Realistic Cell Membranes. Chemical reviews 119, 6184–6226 (2019).

48. T. A. Wassenaar et al., High-Throughput Simulations of Dimer and Trimer Assembly of Membrane Proteins. The DAFT Approach. J Chem Theory Comput 11, 2278–2291 (2015).

49. S. J. Marrink, D. P. Tieleman, Perspective on the Martini model. Chem Soc Rev 42, 6801–6822 (2013).

50. D. H. de Jong et al., Improved Parameters for the Martini Coarse-Grained Protein Force Field. J Chem Theory Comput 9, 687–697 (2013).

51. L. Monticelli et al., The MARTINI Coarse-Grained Force Field: Extension to Proteins. J Chem Theory Comput 4, 819–834 (2008).

52. M. G. Teese, D. Langosch, Role of GxxxG Motifs in Transmembrane Domain Interactions. Biochemistry 54, 5125–5135 (2015).

53. A. Fink, N. Sal-Man, D. Gerber, Y. Shai, Transmembrane domains interactions within the membrane milieu: principles, advances and challenges. Biochimica et biophysica acta 1818, 974–983 (2012).

54. W. P. Russ, D. M. Engelman, The GxxxG motif: a framework for transmembrane helix-helix association. Journal of molecular biology 296, 911–919 (2000).

55. R. F. Walters, W. F. DeGrado, Helix-packing motifs in membrane proteins. Proceedings of the National Academy of Sciences of the United States of America 103, 13658–13663 (2006).

56. S. Li, B. Wu, W. Han, Parametrization of MARTINI for Modeling Hinging Motions in Membrane Proteins. J Phys Chem B 123, 2254–2269 (2019).

57. J. Fantini, F. J. Barrantes, How cholesterol interacts with membrane proteins: an exploration of cholesterol-binding sites including CRAC, CARC, and tilted domains. Front Physiol 4, 31 (2013).

58. R. M. Epand, Cholesterol and the interaction of proteins with membrane domains. Prog Lipid Res 45, 279–294 (2006).

59. I. L. van Genderen, G. J. Godeke, P. J. Rottier, G. van Meer, The phospholipid composition of enveloped viruses depends on the intracellular membrane through which they bud. Biochem Soc Trans 23, 523–526 (1995).

60. G. van Meer, D. R. Voelker, G. W. Feigenson, Membrane lipids: where they are and how they behave. Nature reviews. Molecular cell biology 9, 112–124 (2008).

61. C. Wei et al., HDL-scavenger receptor B type 1 facilitates SARS-CoV-2 entry. Nat Metab 2, 1391–1400 (2020).

62. S. C. Chiliveri, J. M. Louis, R. Ghirlando, J. L. Baber, A. Bax, Tilted, Uninterrupted, Monomeric HIV-1 gp41 Transmembrane Helix from Residual Dipolar Couplings. J Am Chem Soc 140, 34–37 (2018).

63. J. A. Killian, Hydrophobic mismatch between proteins and lipids in membranes. Biochimica et biophysica acta 1376, 401–415 (1998).

64. S. C. Harrison, Viral membrane fusion. Nature structural & molecular biology 15, 690–698 (2008).

65. H. Yao et al., Molecular Architecture of the SARS-CoV-2 Virus. Cell 183, 730–738 e713 (2020).

66. B. Turonova et al., In situ structural analysis of SARS-CoV-2 spike reveals flexibility mediated by three hinges. Science 370, 203–208 (2020).

67. Z. Ke et al., Structures and distributions of SARS-CoV-2 spike proteins on intact virions. Nature 588, 498–502 (2020).

68. F. S. Mesquita et al., S-acylation controls SARS-CoV-2 membrane lipid organization and enhances infectivity. Dev Cell 56, 2790–2807 e2798 (2021).

69. S. Lall, M. K. Mathew, in Membrane Organization and Dynamics, A. Chattopadhyay, Ed. (Springer International Publishing, Cham, 2017), pp. 219–241.

70. A. Senes, M. Gerstein, D. M. Engelman, Statistical analysis of amino acid patterns in transmembrane helices: the GxxxG motif occurs frequently and in association with beta-branched residues at neighboring positions. Journal of molecular biology 296, 921–936 (2000).

71. S. M. Anderson, B. K. Mueller, E. J. Lange, A. Senes, Combination of Calpha-H Hydrogen Bonds and van der Waals Packing Modulates the Stability of GxxxG-Mediated Dimers in Membranes. J Am Chem Soc 139, 15774–15783 (2017).

72. K. R. MacKenzie, J. H. Prestegard, D. M. Engelman, A transmembrane helix dimer: structure and implications. Science 276, 131–133 (1997).

73. E. Li, W. C. Wimley, K. Hristova, Transmembrane helix dimerization: beyond the search for sequence motifs. Biochimica et biophysica acta 1818, 183–193 (2012).

74. P. E. Bragin et al., HER2 Transmembrane Domain Dimerization Coupled with Self-Association of Membrane-Embedded Cytoplasmic Juxtamembrane Regions. Journal of molecular biology 428, 52–61 (2016).

75. T. C. Marcink et al., Intermediates in SARS-CoV-2 spike-mediated cell entry. Sci Adv 8, eabo3153 (2022).

76. S. C. Harrison, Viral membrane fusion. Virology 479-480, 498–507 (2015).

77. T. Tang, M. Bidon, J. A. Jaimes, G. R. Whittaker, S. Daniel, Coronavirus membrane fusion mechanism offers a potential target for antiviral development. Antiviral Res 178, 104792 (2020).

78. G. Griffiths, P. Rottier, Cell biology of viruses that assemble along the biosynthetic pathway. Semin Cell Biol 3, 367–381 (1992).

79. B. Gorai, A. K. Sahoo, A. Srivastava, N. M. Dixit, P. K. Maiti, Concerted Interactions between Multiple gp41 Trimers and the Target Cell Lipidome May Be Required for HIV-1 Entry. J Chem Inf Model 61, 444–454 (2021).

80. R. M. Epand, H. J. Vogel, Diversity of antimicrobial peptides and their mechanisms of action. Biochimica et biophysica acta 1462, 11–28 (1999).

81. C. Subbalakshmi, N. Sitaram, Mechanism of antimicrobial action of indolicidin. FEMS Microbiol Lett 160, 91–96 (1998).

82. H. Jenssen, P. Hamill, R. E. Hancock, Peptide antimicrobial agents. Clin Microbiol Rev 19, 491–511 (2006).

83. D. I. Chan, E. J. Prenner, H. J. Vogel, Tryptophan- and arginine-rich antimicrobial peptides: structures and mechanisms of action. Biochimica et biophysica acta 1758, 1184–1202 (2006).

84. W. Humphrey, A. Dalke, K. Schulten, VMD: visual molecular dynamics. J Mol Graph 14, 33–38, 27-38 (1996).

85. J. Lee et al., CHARMM-GUI Input Generator for NAMD, GROMACS, AMBER, OpenMM, and CHARMM/OpenMM Simulations Using the CHARMM36 Additive Force Field. J Chem Theory Comput 12, 405–413 (2016).

86. E. L. Wu et al., CHARMM-GUI Membrane Builder toward realistic biological membrane simulations. J Comput Chem 35, 1997–2004 (2014).

87. S. Jo, J. B. Lim, J. B. Klauda, W. Im, CHARMM-GUI Membrane Builder for mixed bilayers and its application to yeast membranes. Biophysical journal 97, 50–58 (2009).

88. D. Van Der Spoel et al., GROMACS: fast, flexible, and free. J Comput Chem 26, 1701–1718 (2005).

89. J. Huang, A. D. MacKerell, Jr., CHARMM36 all-atom additive protein force field: validation based on comparison to NMR data. J Comput Chem 34, 2135–2145 (2013).

90. B. Hess, H. Bekker, H. J. C. Berendsen, J. G. E. M. Fraaije, LINCS: A linear constraint solver for molecular simulations. Journal of Computational Chemistry 18, 1463–1472 (1997).

91. T. Darden, D. York, L. Pedersen, Particle mesh Ewald: An N⋅log(N) method for Ewald sums in large systems. The Journal of chemical physics 98, 10089–10092 (1993).

92. H. J. C. Berendsen, J. P. M. Postma, W. F. Vangunsteren, A. Dinola, J. R. Haak, Molecular-Dynamics with Coupling to an External Bath. Journal of Chemical Physics 81, 3684–3690 (1984).

93. W. G. Hoover, Canonical dynamics: Equilibrium phase-space distributions. Phys Rev A Gen Phys 31, 1695–1697 (1985).

94. S. Nosé, A molecular dynamics method for simulations in the canonical ensemble. Molecular Physics 52, 255–268 (1984).

95. M. Parrinello, A. Rahman, Polymorphic Transitions in Single-Crystals - a New Molecular-Dynamics Method. Journal of Applied Physics 52, 7182–7190 (1981).

96. W. J. Allen, J. A. Lemkul, D. R. Bevan, GridMAT-MD: a grid-based membrane analysis tool for use with molecular dynamics. J Comput Chem 30, 1952–1958 (2009).

97. S. J. Marrink, H. J. Risselada, S. Yefimov, D. P. Tieleman, A. H. de Vries, The MARTINI force field: coarse grained model for biomolecular simulations. J Phys Chem B 111, 7812–7824 (2007).

98. G. Bussi, D. Donadio, M. Parrinello, Canonical sampling through velocity rescaling. The Journal of chemical physics 126, 014101 (2007).

99. W. Song et al., PyLipID: A Python Package for Analysis of Protein-Lipid Interactions from Molecular Dynamics Simulations. J Chem Theory Comput 18, 1188–1201 (2022).

100. T. A. Wassenaar, K. Pluhackova, R. A. Bockmann, S. J. Marrink, D. P. Tieleman, Going Backward: A Flexible Geometric Approach to Reverse Transformation from Coarse Grained to Atomistic Models. J Chem Theory Comput 10, 676–690 (2014).

