## Supplementary material for "Sequence of the SARS-CoV-2 spike transmembrane domain makes it inherently dynamic"

### for

**Table S1. Summary of simulations**

| Description | Sequence | Box size ($nm^3$) | Type [a] | Replicates x Duration | Simulated Time [b] |
|---|---|---|---|---|---|
| TMH in PC | $S_{1196}$–$S_{1249}$ | 8 x 8 x 14 | AA | 3 x 0.5 μs | 1.5 μs |
| PC | – | 8 x 8 x 8.2 | AA | 3 x 0.1 μs | 0.3 μs |
| T1 (3 helices in PC) | $W_{1212}$–$C_{1236}$ | 11.8 x 10.1 x 8.4 | CG | 100 x 4 μs | 400 μs |
| T2 (3 helices in PC) | $Y_{1215}$–$C_{1241}$ | 13.5 x 11.7 x 10 | CG | 100 x 4 μs | 400 μs |
| T3 (3 helices in PC) | $K_{1211}$–$M_{1237}$ | 16 x 13.2 x 10 | CG | 100 x 4 μs | 400 μs |
| T1 + 30% Chol | Same as T1 | 13.1 x 11.2 x 9.5 | CG | 50 x 4 μs | 200 μs |
| T2 + 30% Chol | Same as T2 | 13.2 x 11.5 x 10 | CG | 50 x 4 μs | 200 μs |
| T3 + 30% Chol | Same as T3 | 15.8 x 13.8 x 10 | CG | 50 x 4 μs | 200 μs |
| D1 (2 helices in PC) | $W_{1214}$–$C_{1236}$ | 12.3 x 10.6 x 9.3 | CG | 75 x 1.5 μs | 112.5 μs |
| D2 (2 helices in PC) | $W_{1212}$–$L_{1234}$ | 12.3 x 10.6 x 9.3 | CG | 75 x 1.5 μs | 112.5 μs |

[a] Refers to the nature of simulation. AA: all-atom (CHARMM36); CG: coarse-grained (Martini)

[b] Aggregated simulation time

Box size refers to the starting dimensions of the simulation box

PC refers to POPC (1-palmitoyl-2-oleoyl-glycerophosphatidylcholine) bilayer

Chol refers to cholesterol in mol%

#### SUPPLEMENTARY FIGURES

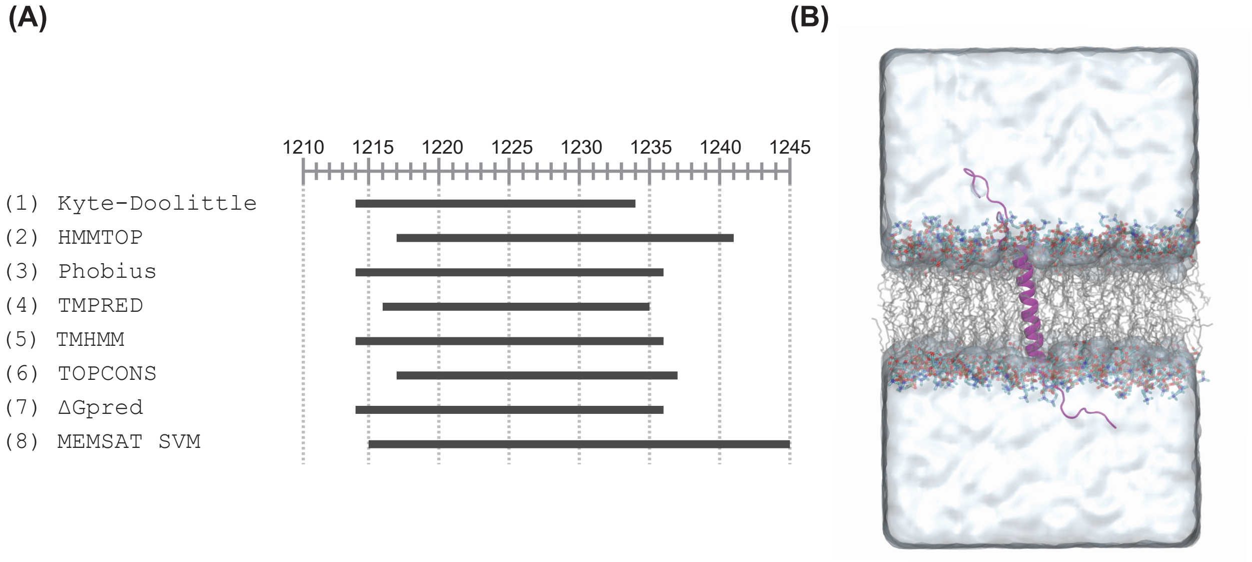


**Figure S1: Prediction of the transmembrane (TM) helix in the S2 domain of the SARS-CoV-2 spike protomer.** (**A**) Residue numbering shown at the top follows the numbering in full length spike. Horizontal black lines mark the TM helix residues predicted by different algorithms (numbered 1-8). The ExPASy server implementation of the Kyte-Doolittle algorithm was used (https://web.expasy.org/protscale/). The other algorithms were used from their native webservers with default settings. (**B**) Starting conformation for the atomistic simulations. A hydrated POPC membrane-embedded 54-residue long polypeptide (Ser1196 to Ser1249) comprising the helical Pro1213 to Met1237 (protein, purple; lipid acyl chains, gray; lipid headgroups colored by identity of the atoms (Oxygen, red; Carbon, cyan and Nitrogen, blue); water in surface view; Hydrogen atoms of the lipid molecules are omitted for clarity).

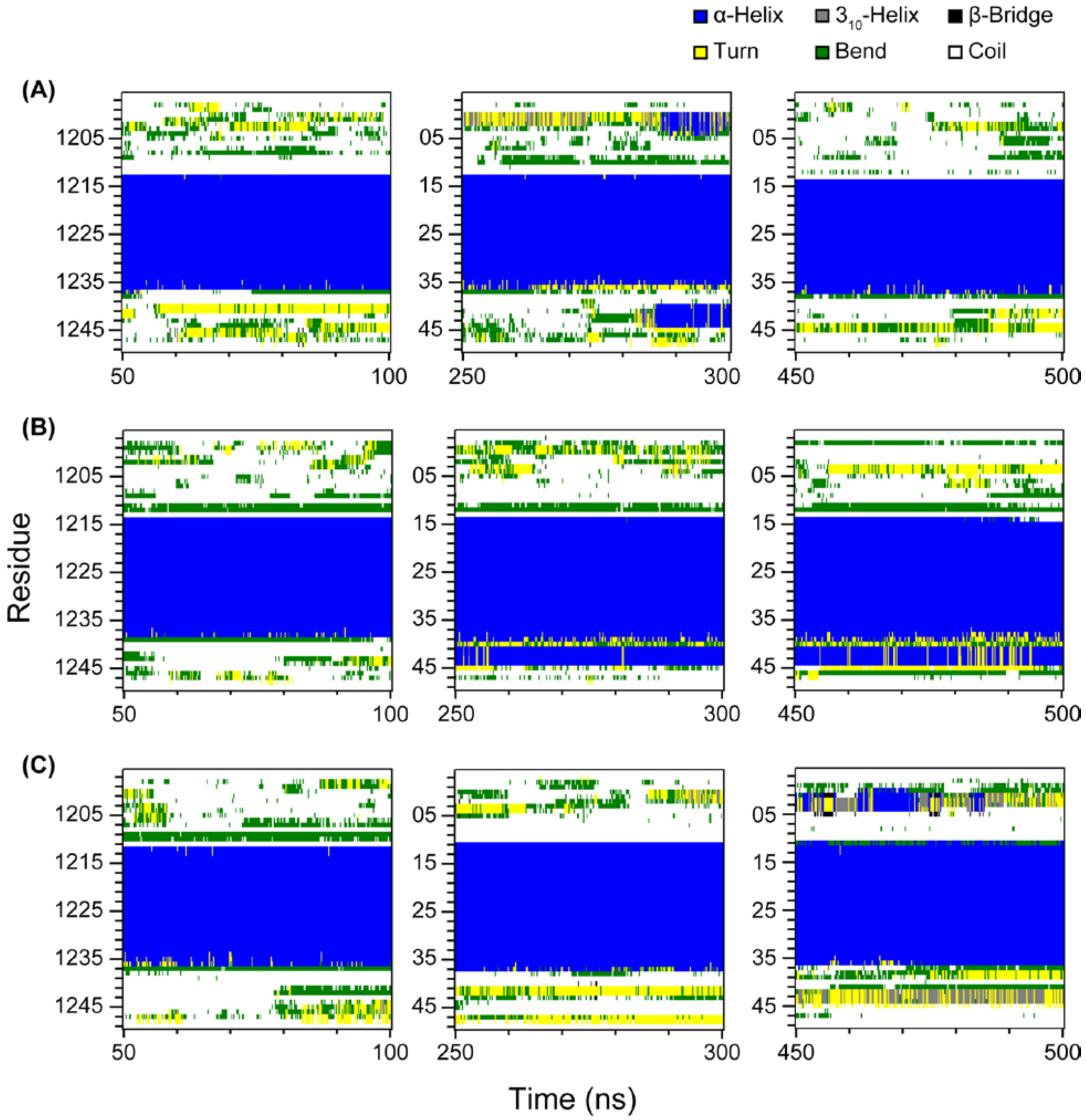


**Figure S2: Secondary structure analysis of the atomistic simulations of the spike protomer segment, Ser1196 to Ser1249.** (**A-C**) Results from secondary structure analysis of the three replicate atomistic trajectories of 500 ns each. Three windows (left to right) of 50 ns each starting at 50, 250 and 450 ns for each trajectory are shown. Each point in these plots corresponds to the frame-wise progression of per residue secondary structure estimated by DSSP. Different secondary structure elements are colored as per the legends above the figure.

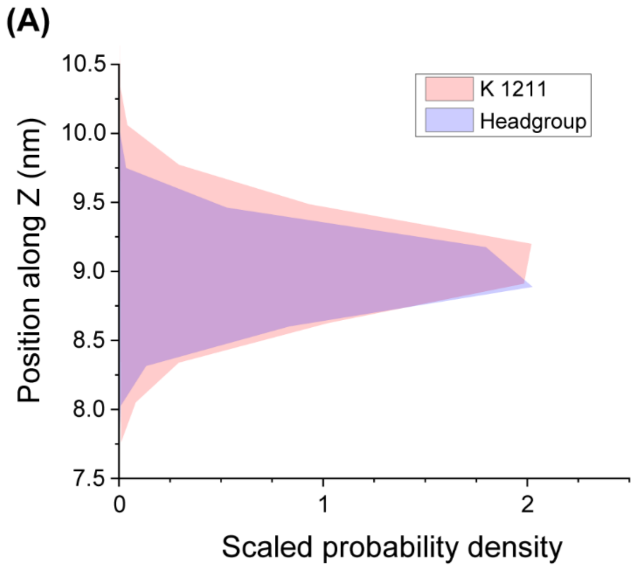


**Figure S3**: **Perpendicular distance of Lys1211 from the membrane plane.** Scaled probability density of the Cα atom of Lys1211 compared with that of the headgroup of the relevant membrane leaflet emphasizes the snorkeling by the Lys residue with excursions into water. For this analysis, the three atomistic replicate trajectories were combined and the bilayer was centered with respect to the simulation box.

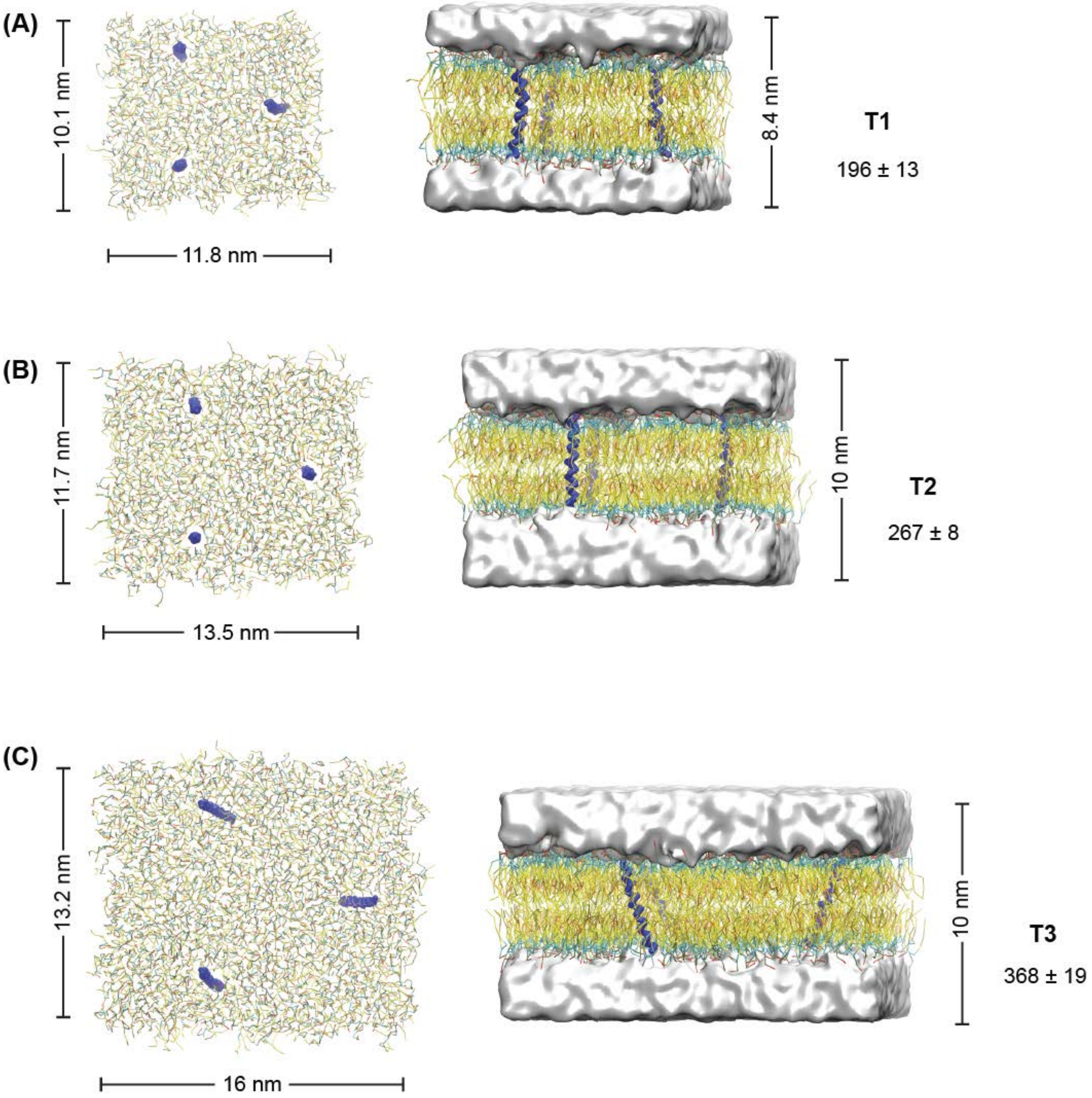


**Figure S4: Representative starting structures of the simulations.** The top (left) and side (right) view of the simulation box used in (**A**) T1, (**B**) T2 and (**C**) T3 simulations. (three identical helices per simulation, blue trace; lipid headgroup, green lines, lipid acyl chains, yellow and orange lines; water, white surface; water removed from top view for clarity). The x, y, and z dimensions are listed. Number on the right is the mean ± SD of lipids per leaflet (n=15).

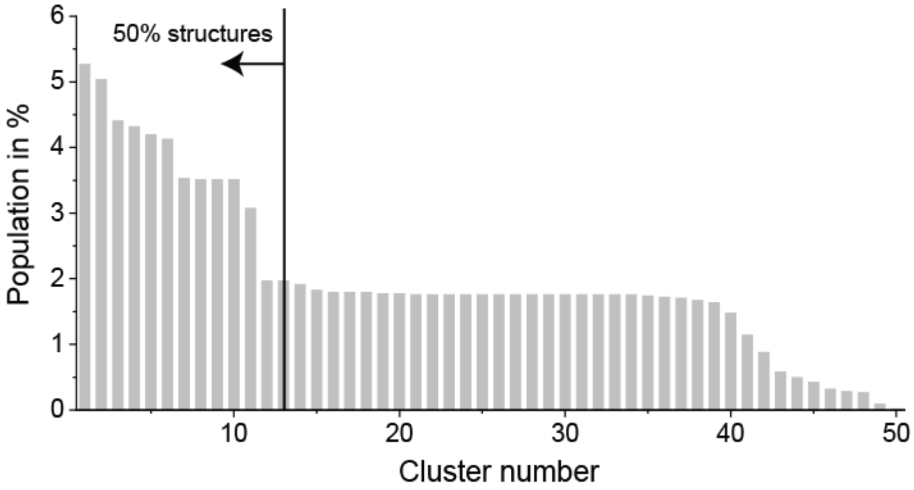


**Figure S5: Population of conformations observed for all the T1 trimerized trajectories.** 50 clusters differing at 5 Å RMSD were observed from the last 50 ns combined from 53 trajectories, each, that ended trimeric. The population distribution of the clusters shows that the top 13 clusters contribute 50% of all the trimeric conformations sampled. The top five clusters are discussed here.

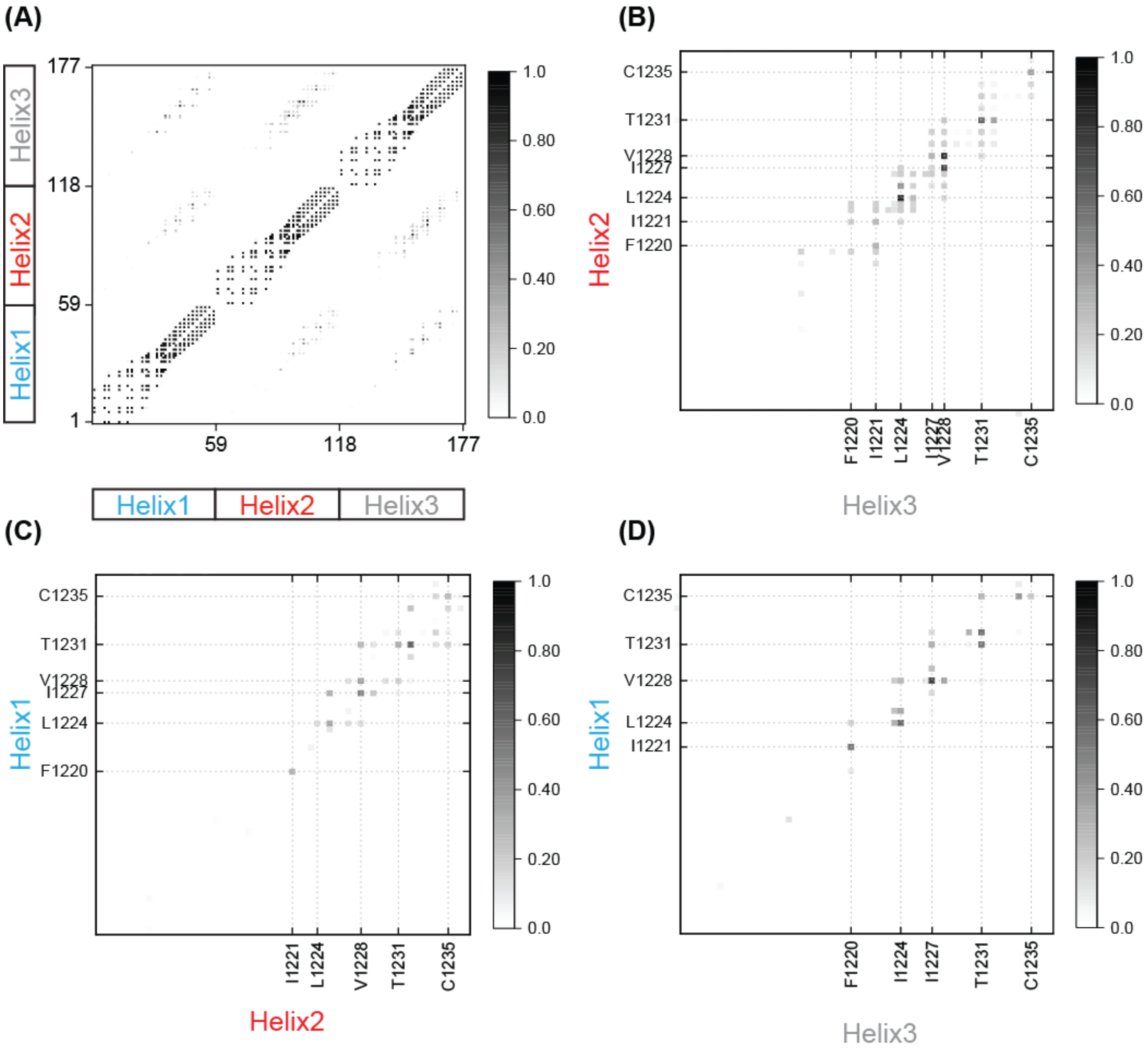


**Figure S6: Contact map of the symmetric trimer shown in Fig. 3C.** (**A**) Bead vs bead contact map of the symmetric T1 trimer. Darker colors signify a greater observed frequency of two beads (coarse-grained particles) being within 5 Å of one another. A value of 1 in the color bars means that the contact was found in all structures in that cluster. (**B-D**) Zoomed views from (**A**) the three pairs of inter-helix contacts formed between the three T1 helices that undergo trimerization show that they are similar. The absence of contacts between the TRACS means that they do not interact with one another.

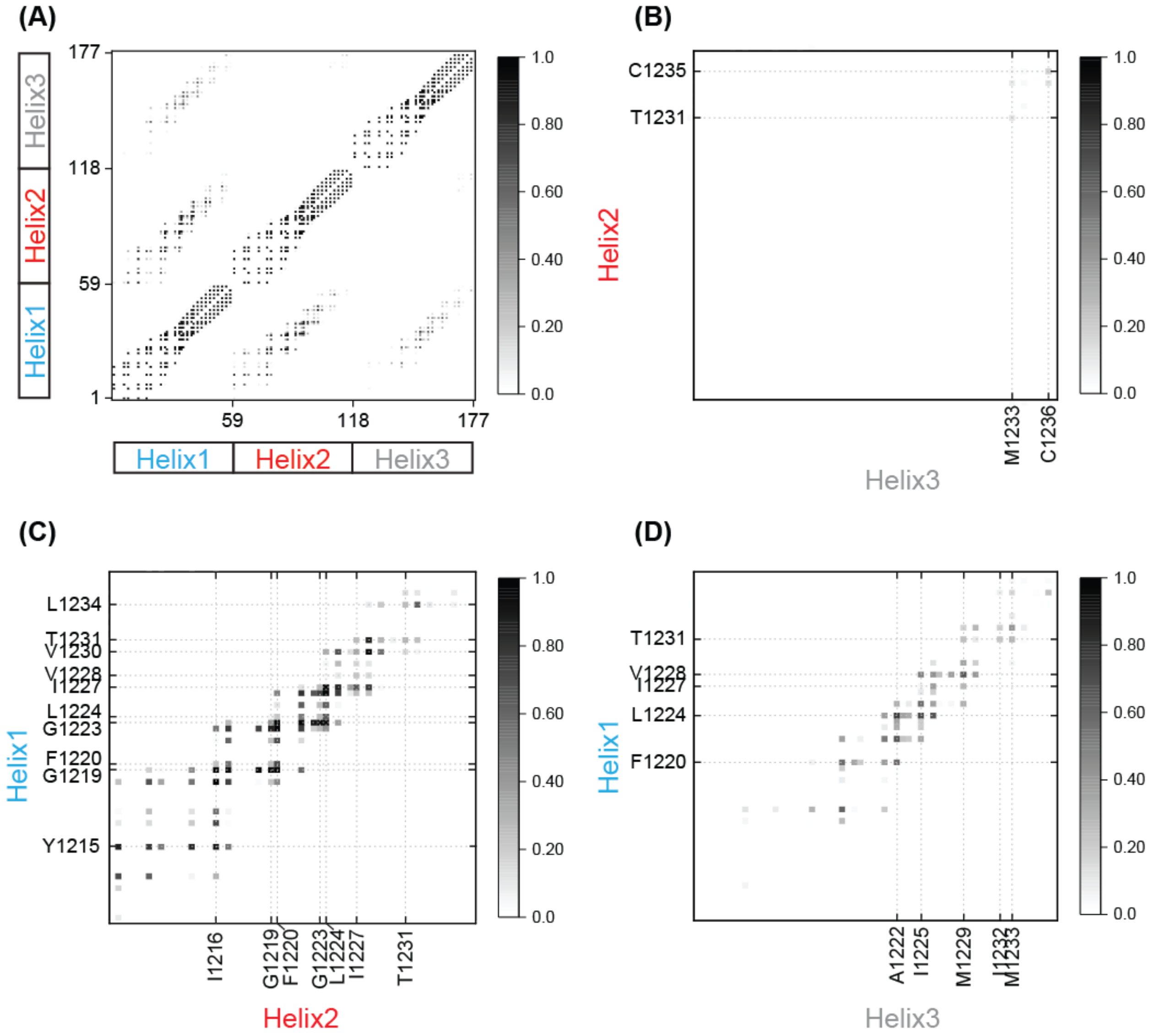


**Figure S7: Contact map of the asymmetric trimer shown in Fig. 3D.** Bead vs bead contact map of the asymmetric T1 trimer calculated in a manner similar to **Fig. S6**. Helix 2 and Helix 3 barely interact with each other.

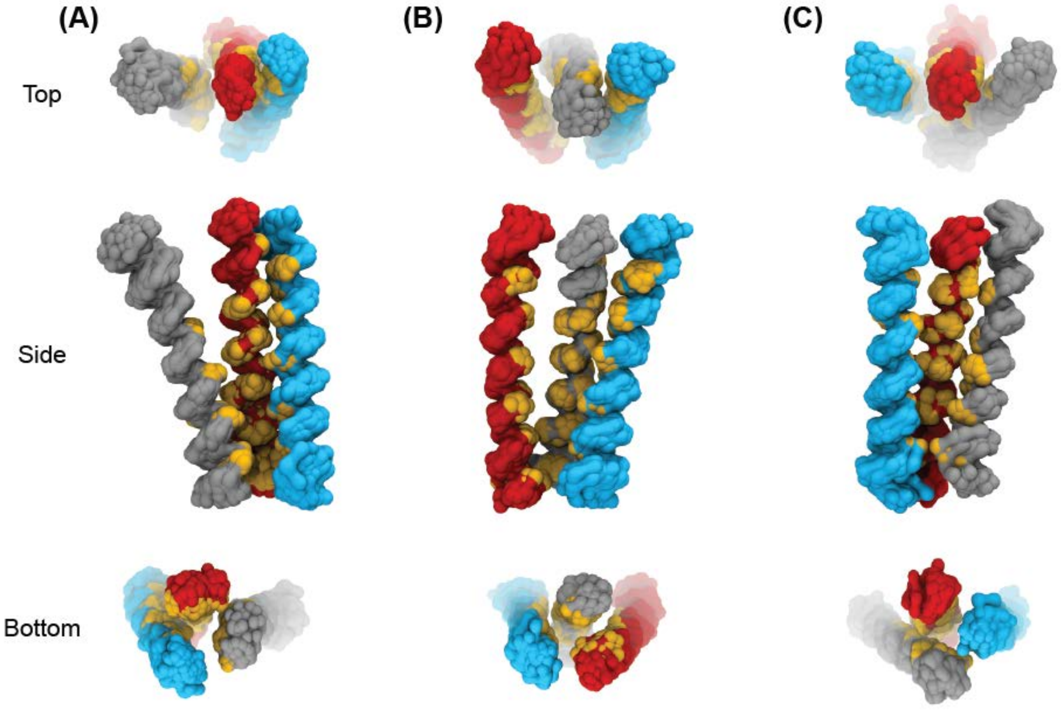


**Figure S8: Other high occupancy structural clusters from T1 simulations.** Three clusters of T1 conformations, each showing three T1 helices colored red, cyan and gray (interacting residues in yellow). The N-terminus is at the top. The C-terminus (down) is more compact and approaching three-fold symmetry while the N-terminus (up) is splayed.

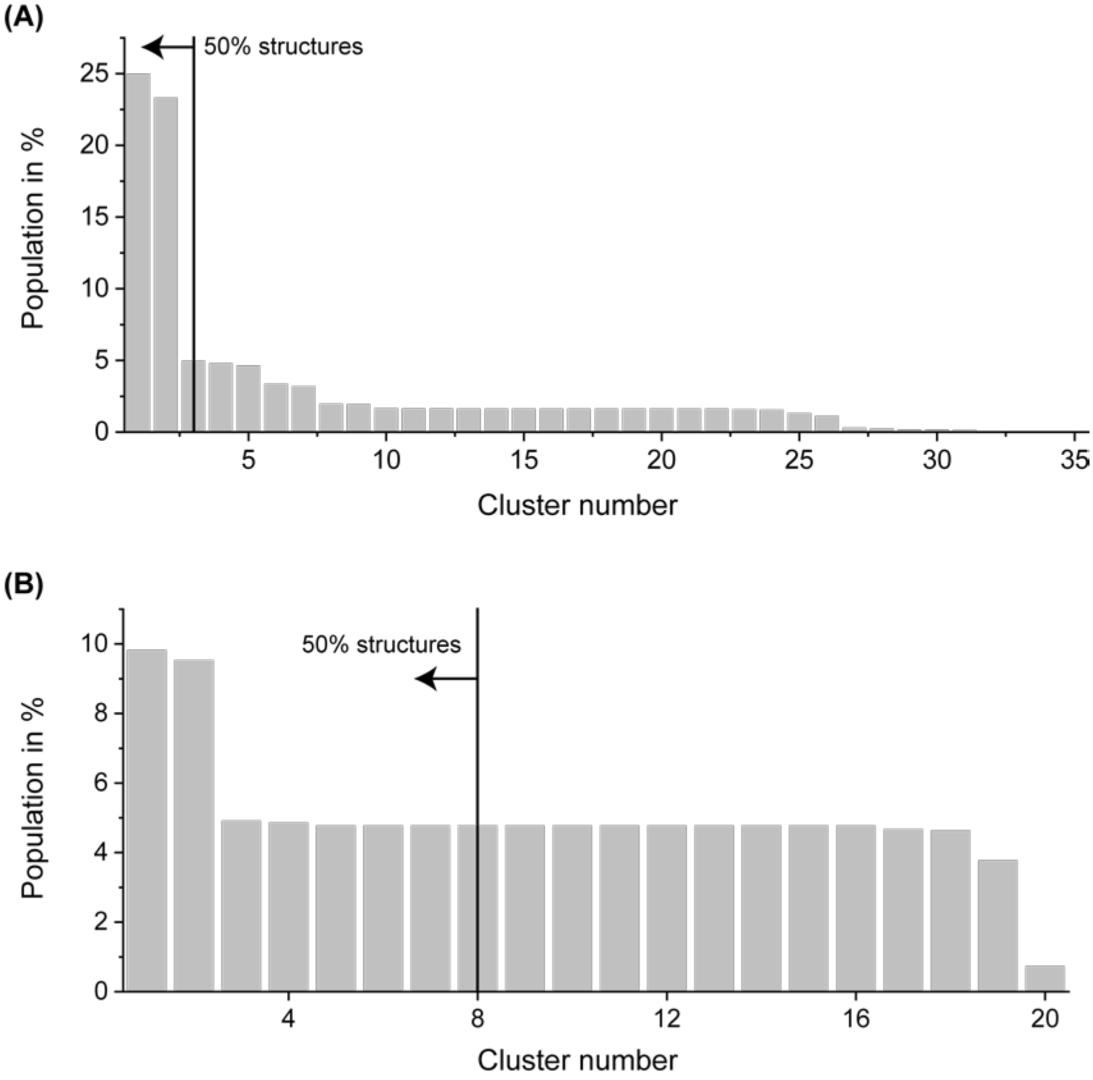


**Figure S9**: **Population of clusters of conformations observed in the last 50 ns of all the (A) T2 and (B) T3 trimerized trajectories.** 35 clusters differing at 5 Å RMSD were observed in T2 and 20 clusters were observed in T3. The three most populated clusters make up >50% of all the trimeric conformations in T2. Whereas, the T3 conformational ensemble was highly diverse as the 21 trimerized trajectories classified into 20 conformation clusters. Cluster 1 from T2 is shown in **Fig. 4D**.

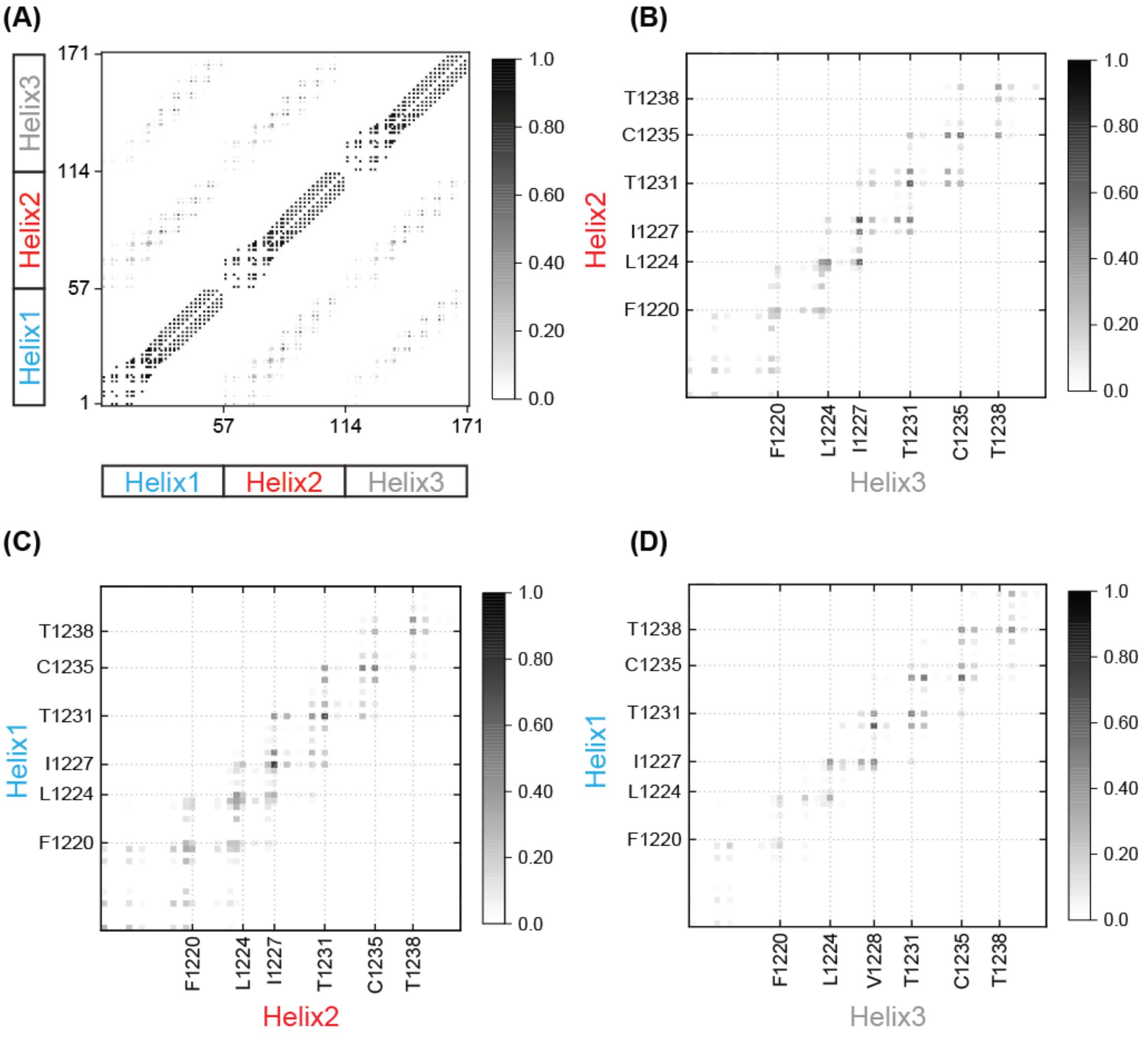


**Figure S10: Contact map of the T2 symmetric trimer from Fig. 4D.** Bead vs bead contact map of the symmetric T2 trimer. T2 lacks two of the N-terminal TRACS residues which did not show homotypic interactions in T1 (**Fig. S6**).

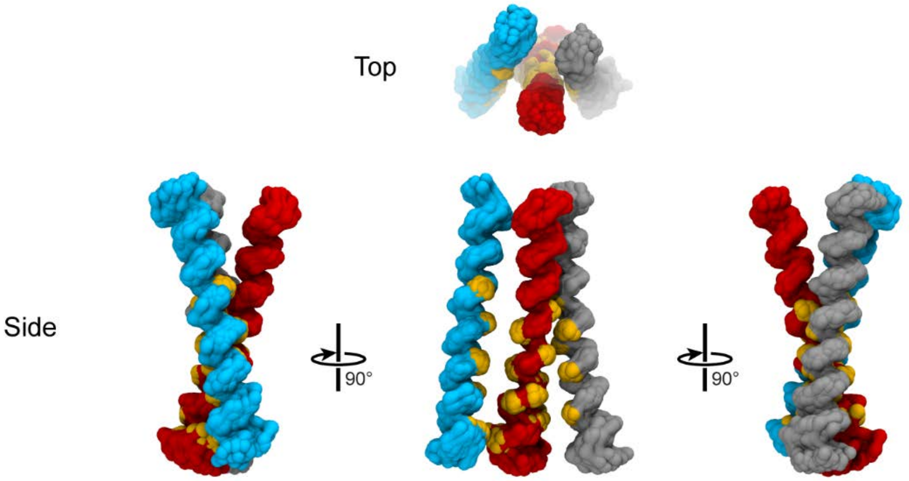


**Figure S11**: **The most populated T3 trimer.** Major cluster showing the three T3 helices colored red, cyan and gray (interacting residues in yellow). The structure can be interpreted as a left-handed dimer and a right-handed dimer joined at a common central helix (red). The N-terminus is at the top.

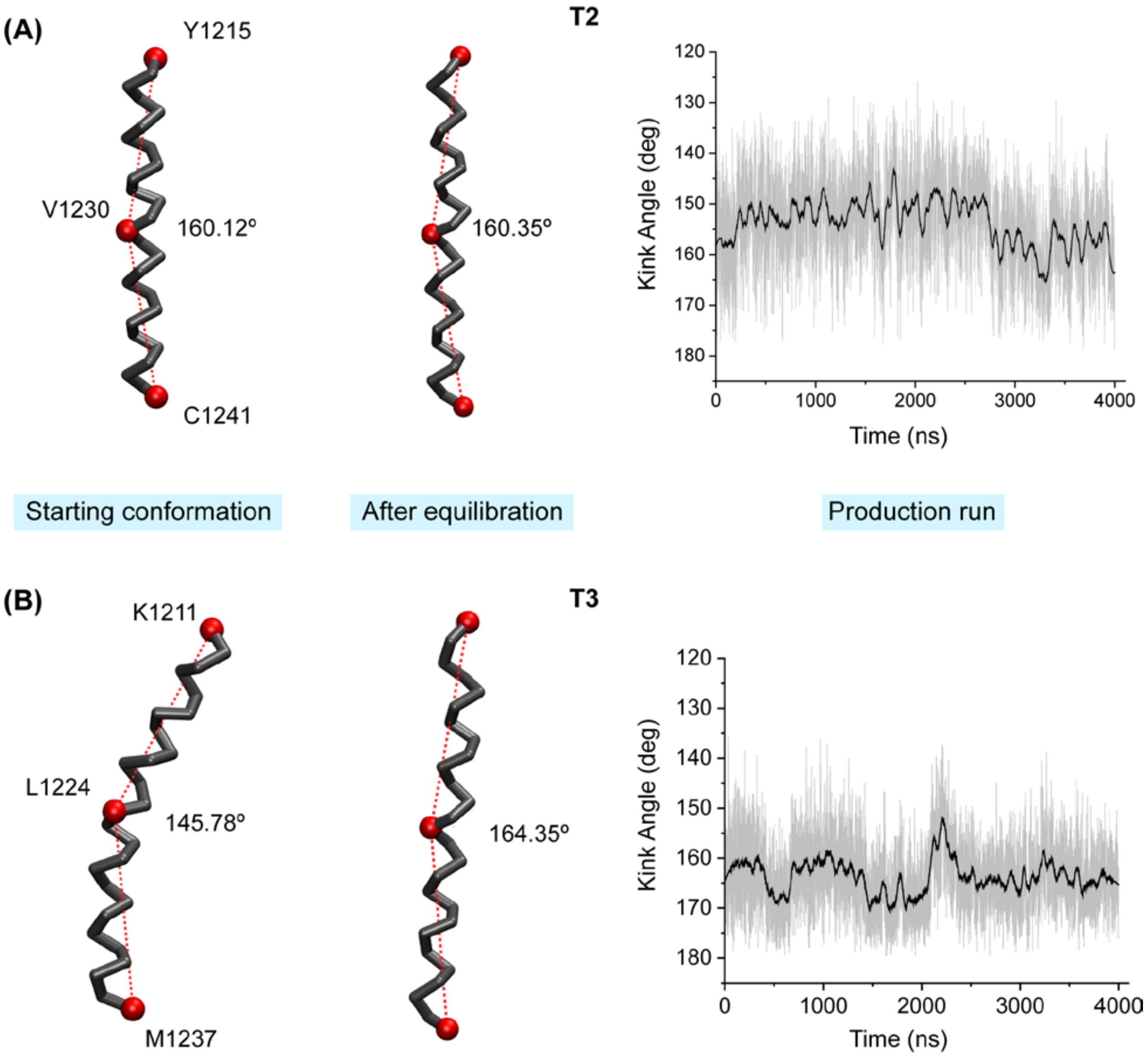


**Figure S12: Deviation of helical bend in the Martini force field.** The coarse-grained helical models of (**A**) T2 and (**B**) T3 are shown as a backbone trace. The three red spheres indicate Cα beads of the residues at the beginning, middle and end of the construct and the angle formed by the three beads is listed. The angle changes even during the equilibration phase of the simulation. Also the same angle is tracked along a production run (right, simulation data in gray and 400 point average overlaid in black). Note the initial difference in angle between the starting T2 (upper left) and T3 (lower left) conformations, and its straightening during the equilibration phase.

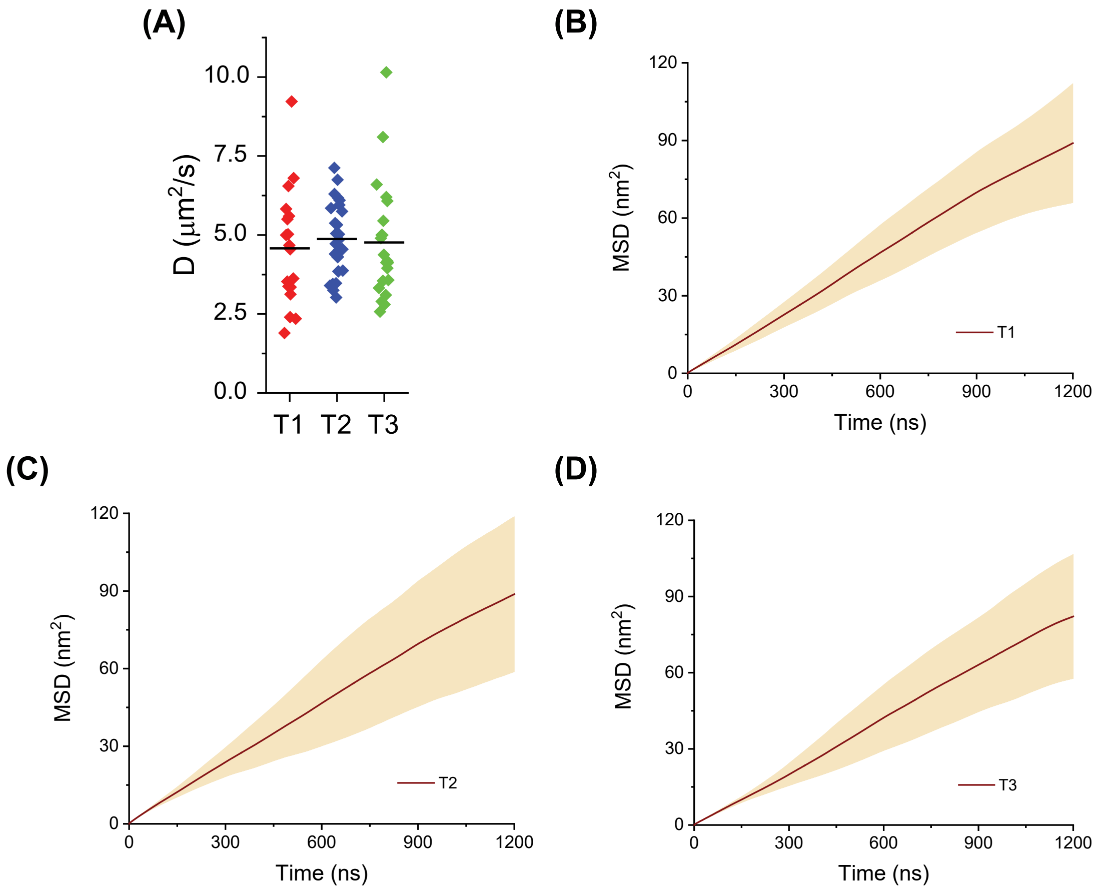


**Figure S13: Diffusion coefficient (D) for the TM helices in POPC bilayers.** (**A**) Each data point indicates the estimated diffusion coefficient for a helix in the POPC bilayer from one simulation. The values were calculated by fitting straight lines to the mean square displacement (MSD) vs time plots over the first 1.2 μs of trajectories. The horizontal line marks the mean value. (**B-D**). Mean (brown line) ± SD (shaded region) of multiple helices from more than 10 independent replicate simulations used to calculate the data points in (**A**).

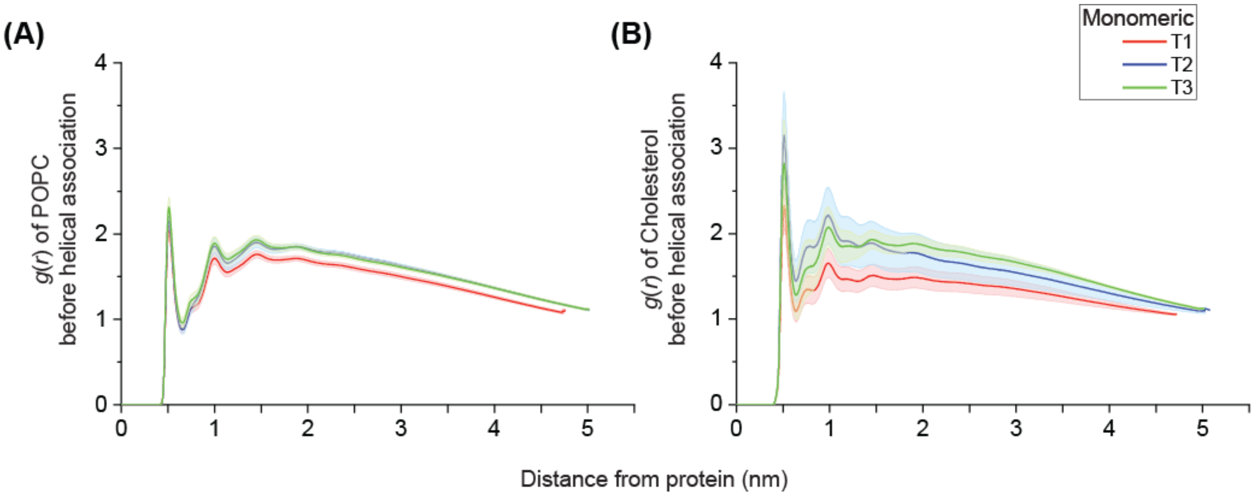


**Figure S14: Radial distribution function** (RDF; *g*(r)) of (**A**) POPC and (**B**) cholesterol (chol) around monomeric helices for the T1 (red), T2 (blue) and T3 (green) simulations. The RDFs were calculated using the initial part of the simulation trajectories before the first association event occurred. (**A**) POPC simulations and (**B**) POPC:chol bilayers. Mean (colored line) ± SD (shaded region) from more than 10 independent replicate simulations are shown. **Fig. 5B** shows a cropped version of **Fig. S14B** without the SD.

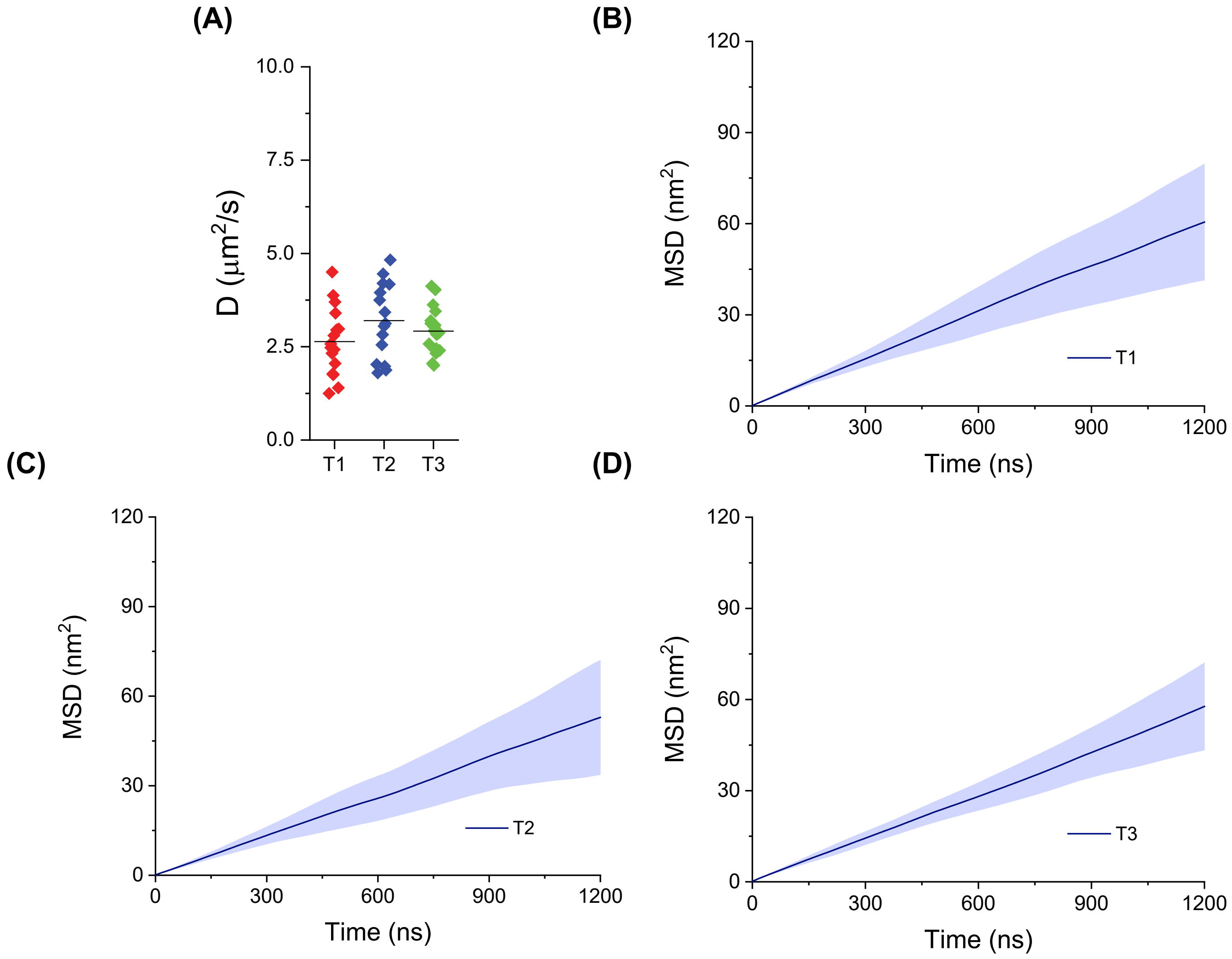


**Figure S15**: **Diffusion coefficient (D) for the TM helices in POPC:chol bilayer.** (**A**) Each data point indicates the estimated diffusion coefficient for a helix in the POPC:chol bilayer in one simulation. Compare with **Fig. S13.** (**B-D**). Mean (blue line) ± SD (shaded region) of multiple helices from more than 10 independent replicate simulations used to calculate the data points in (**A**).

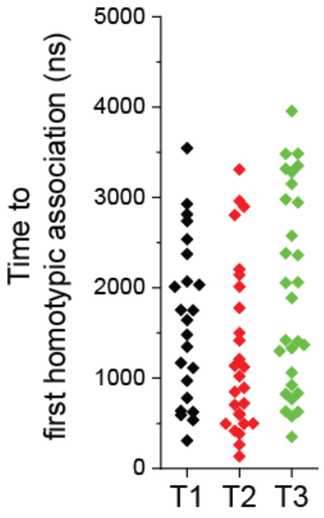


**Figure S16: Time taken for the first inter-helix association event in POPC:chol bilayer.** Each data point indicates the time before the first helix-helix association in one simulation. Data from more than 20 trimerizing simulations for T1, T2 and T3.

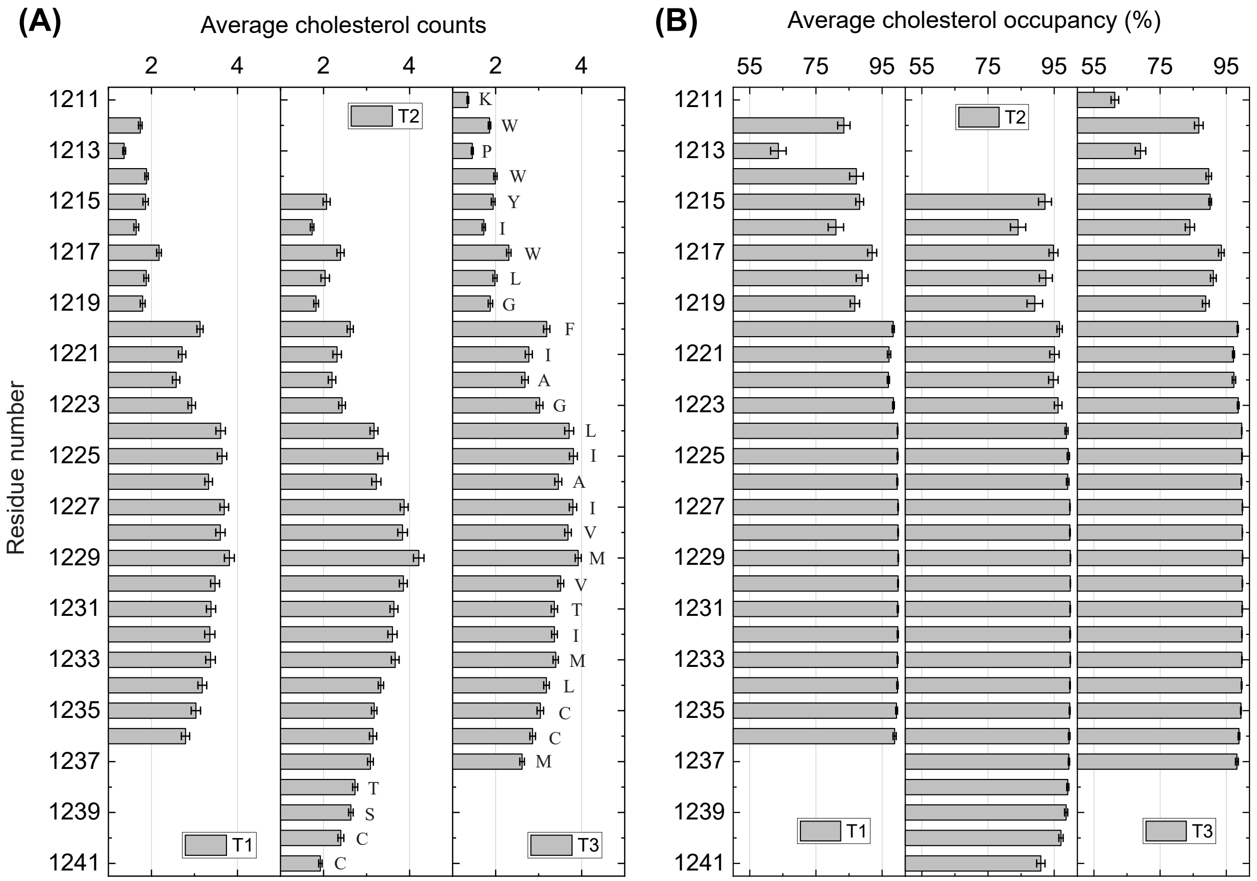


**Figure S17: Interaction of chol with monomeric TM helices.** (**A**) Average number of chol molecules observed within 5.5 Å to 10 Å of the monomeric TM helices. (**B**) Average time occupancy for chol molecules near a particular residue of a monomeric helix. Mean ± SD from 14 independent replicate simulations. Related to **Fig. 5C**.

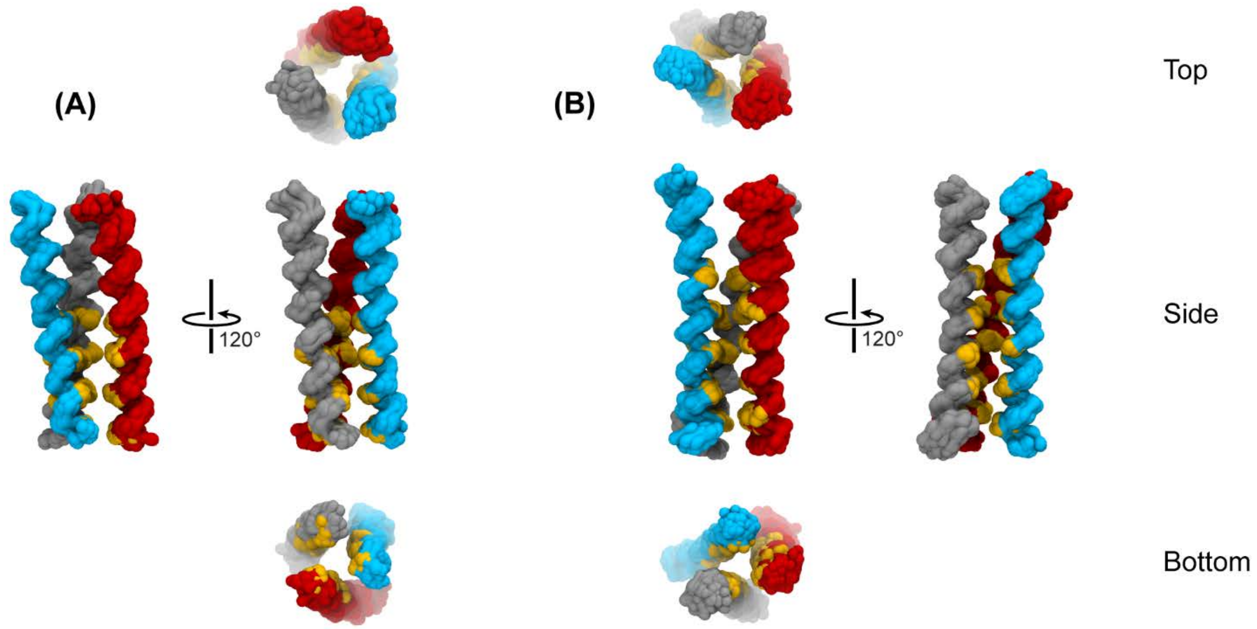


**Figure S18: The most populated cluster from trimerized simulations in POPC:chol bilayers.** Symmetric (**A**) T1 trimer (**B**) T2 trimer conformations. Helices colored red, cyan and gray (backbone of interacting residues in yellow). Top refers to the N-terminus. Compare with **Figs. 3A** and **4D**, respectively.

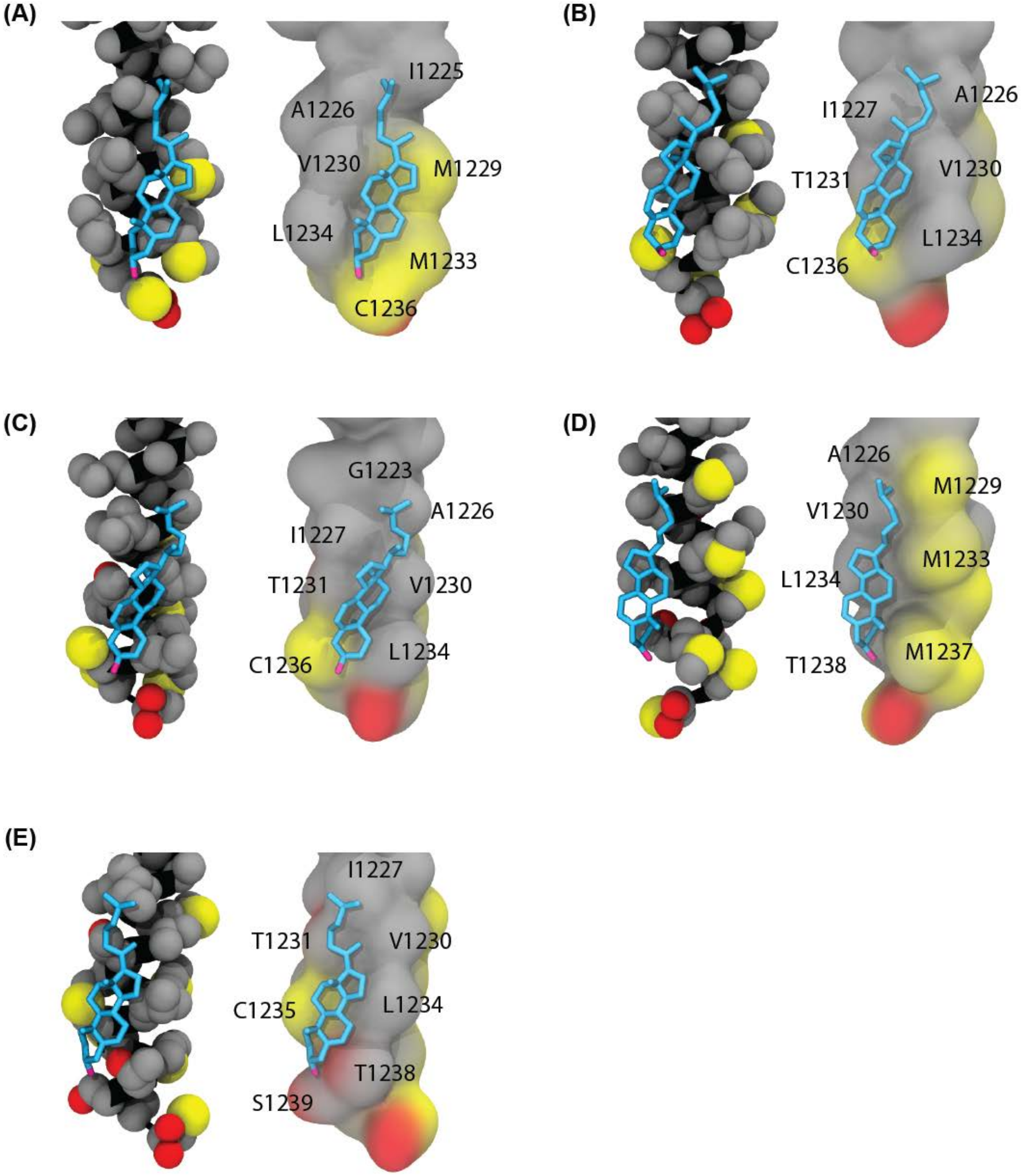


**Figure S19: Frequently encountered poses of chol** (Carbon, cyan; Oxygen, magenta) **bound to the C-terminal half of a monomeric TM helix (A-E)** from backmapped structures (Backbone, black; side chain Carbon, gray; Oxygen, red; Sulphur, yellow; Hydrogens omitted for clarity). Two views of the helix are shown side by side (space filling, left and surface, right) to highlight the extensive packing between the chol molecule and the TM helix. The hydroxyl group of chol is found interacting with the phosphatidylcholine lipid headgroups (not shown).

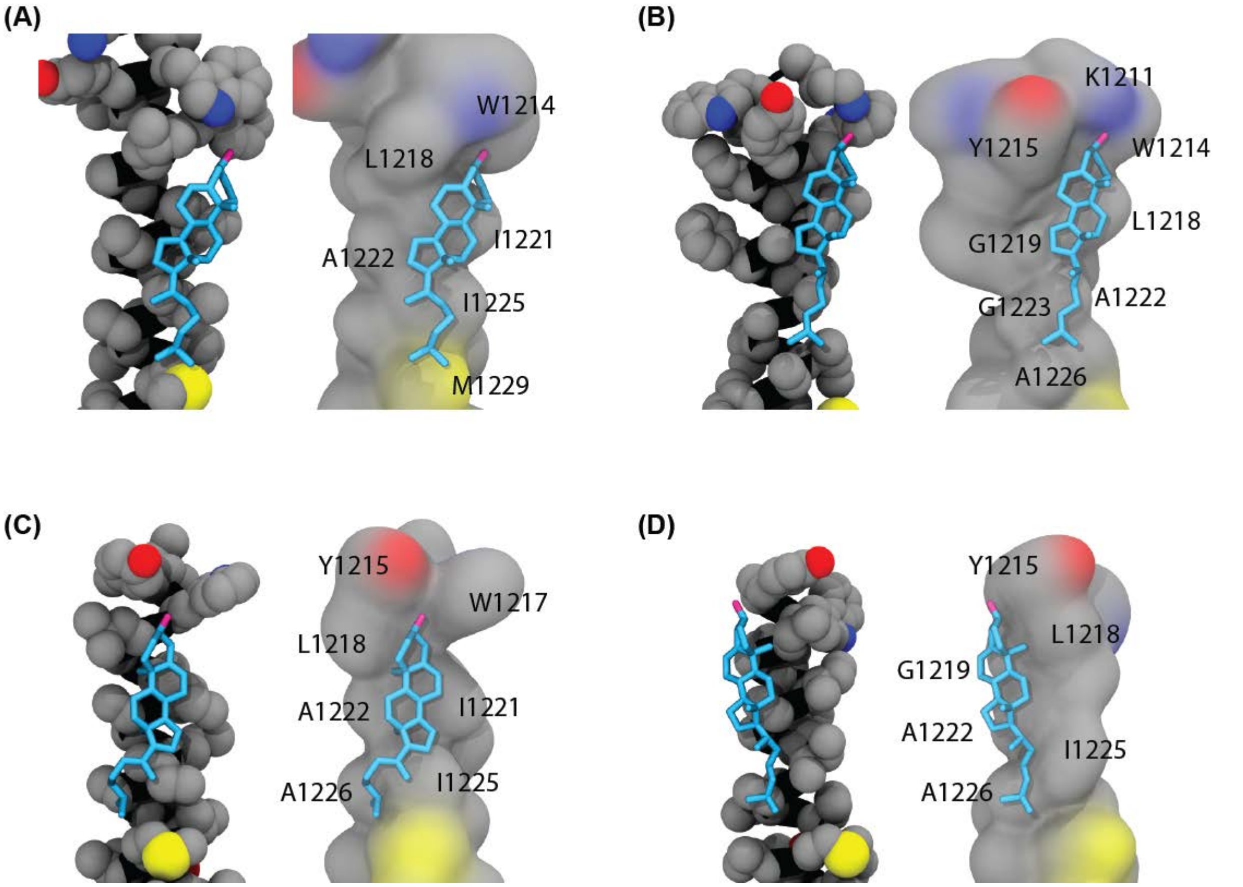


**Figure S20: Frequently encountered poses of chol** (Carbon, cyan; Oxygen, Magenta) **bound to the N-terminal half of a monomeric TM helix (A-D)** from backmapped structures (Backbone, black; side chain Carbon, gray; Oxygen, red; Nitrogen, blue; Sulphur, yellow; Hydrogens omitted for clarity). Two views of the helix are shown side by side (space filling, left and surface, right) to highlight the extensive packing between the chol molecule and the TM helix. The hydroxyl group of chol is found hydrogen bonded to either the indole nitrogen on one of the Trp sidechains or the phosphatidylcholine headgroup (not shown). These poses are rarer compared to those seen for chol bound to the C-terminus in **Fig. S19**.

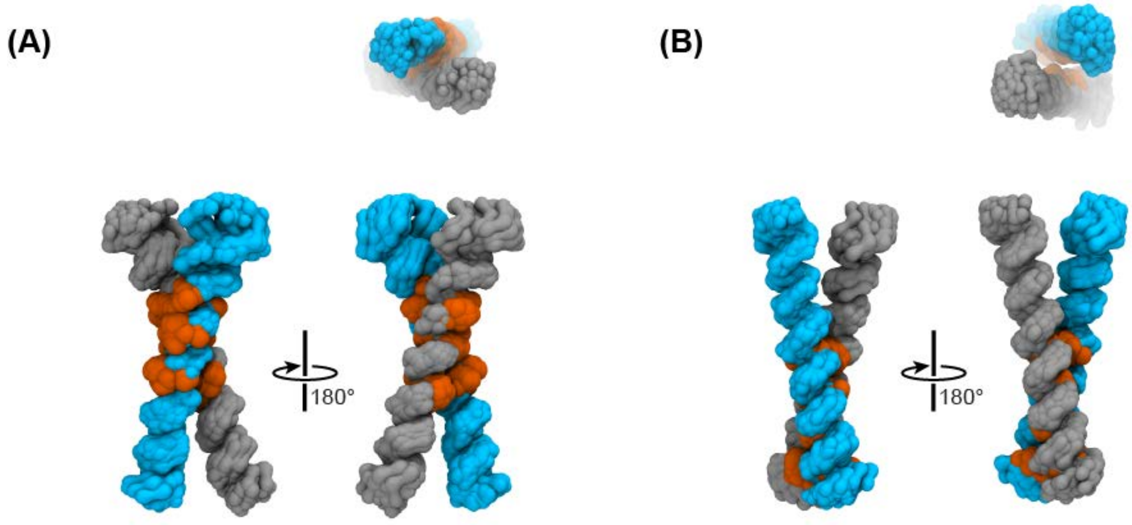


**Figure S21: Two of the highest populated structural clusters from those T3 trimerizing simulations which ended as dimers.** These are comparable to the structures obtained from the dimerizing simulations (**Fig. 7**). (**A**) GxxxG mediated right handed dimers (**Fig. 7C**) and (**B**) left-handed dimers (**Fig. 7D**). (Helical backbones in cyan and gray, backbones of the interacting residues in orange; view looking down from the N-terminus separately shown on top).
